# Biology and physics of heterochromatin-*like* domains/complexes

**DOI:** 10.1101/2020.07.19.210518

**Authors:** Prim B. Singh, Stepan N. Belyakin, Petr P. Laktionov

## Abstract

The hallmarks of constitutive heterochromatin, HP1 and H3K9me2/3, assemble heterochromatin-*like* domains/complexes *outside* canonical constitutively heterochromatic territories where they regulate chromatin-templated processes. Domains are more than 100kb in size; complexes less than 100kb. They are present in the genomes of organisms ranging from fission yeast to man, with an expansion in size and number in mammals. Some of the likely functions of the domains/complexes include silencing of the donor mating type region in fission yeast, regulation of mammalian imprinted genes and the phylotypic progression during vertebrate development. Far *cis*- and *trans*-contacts between micro-phase separated domains/complexes in mammalian nuclei contribute to the emergence of epigenetic compartmental domains (ECDs) detected in Hi-C maps. We speculate that a thermodynamic description of micro-phase separation of heterochromatin-*like* domains/complexes will require a gestalt shift away from the monomer as the “*unit of incompatibility*”, where it is the choice of monomer that determines the sign and magnitude of the Flory-Huggins parameter, χ. Instead, a more dynamic structure, the oligo-nucleosomal “clutch”, consisting of between 2 to 10 nucleosomes is both the long sought-after secondary structure of chromatin and its unit of incompatibility. Based on this assumption we present a simple theoretical framework that enables an estimation of χ for domains/complexes flanked by euchromatin and thereby an indication of their tendency to phase separate. The degree of phase separation is specified by χN, where N is the number of “clutches” in a domain/complex. Our approach may provide an additional tool for understanding the biophysics of the 3D genome.

## 1. Introduction

There is an intimate relationship between gene regulation, chromatin structure and genome organisation [1]. The kernel from which our understanding of this relationship grew can be found in studies on constitutive heterochromatin, especially with the phenomenon of position-effect variegation (PEV) in *Drosophila* (for reviews see [2–5]). Nine decades of work on PEV concluded that changing the position of a gene with respect to the heterochromatin-euchromatin boundary can affect its chromatin structure and that, in turn, affects its expression as manifest by phenotypic variegation and changes in transcription [6–10]. Saturation mutagenesis screens identified modifiers of PEV and molecular characterisation of their wild-type (wt) gene products showed they encode structural components and enzymatic activities that regulate the assembly of heterochromatin [5, 11–13]. Two of the modifiers encode Heterochromatin Protein 1 (HP1) and H3K9HMTases that generate the H3K9me2/3^1^ histone modification that is bound by the chromo domain (CD) of HP1 [15] (Figures 1A and B). HP1 and the H3K9me3 modification are highly conserved across eukaryotes and represent hallmarks of constitutive heterochromatin [16,17] that are enriched at constitutively heterochromatic chromosomal territories of nearly all eukaryotic chromosomes. These territories include peri-centric heterochromatin surrounding the centromeres, (sub-) telomeric and (peri-) nucleolar organiser regions (NORs), with both hallmarks being found at these sites in organisms as distantly related as fission yeast through *Drosophila* to man [18–25]. Notable exceptions are the chromosomes of budding yeast where the silent information regulator (Sir) complex is assembled at heterochromatic territories (telomeres and NORs) by establishing and recognizing a pattern of de-acetylated histones, especially hypo-acetylated H4K16 [26]. Colocalisation of HP1 and H3K9me3 to constitutive heterochromatin arose early in the evolution of eukaryotes with the common ancestor of fission yeast and man living around one billion years ago [27]; budding and fission yeasts diverged from each other at around the same time both did from man [28].

**Figure 1:**
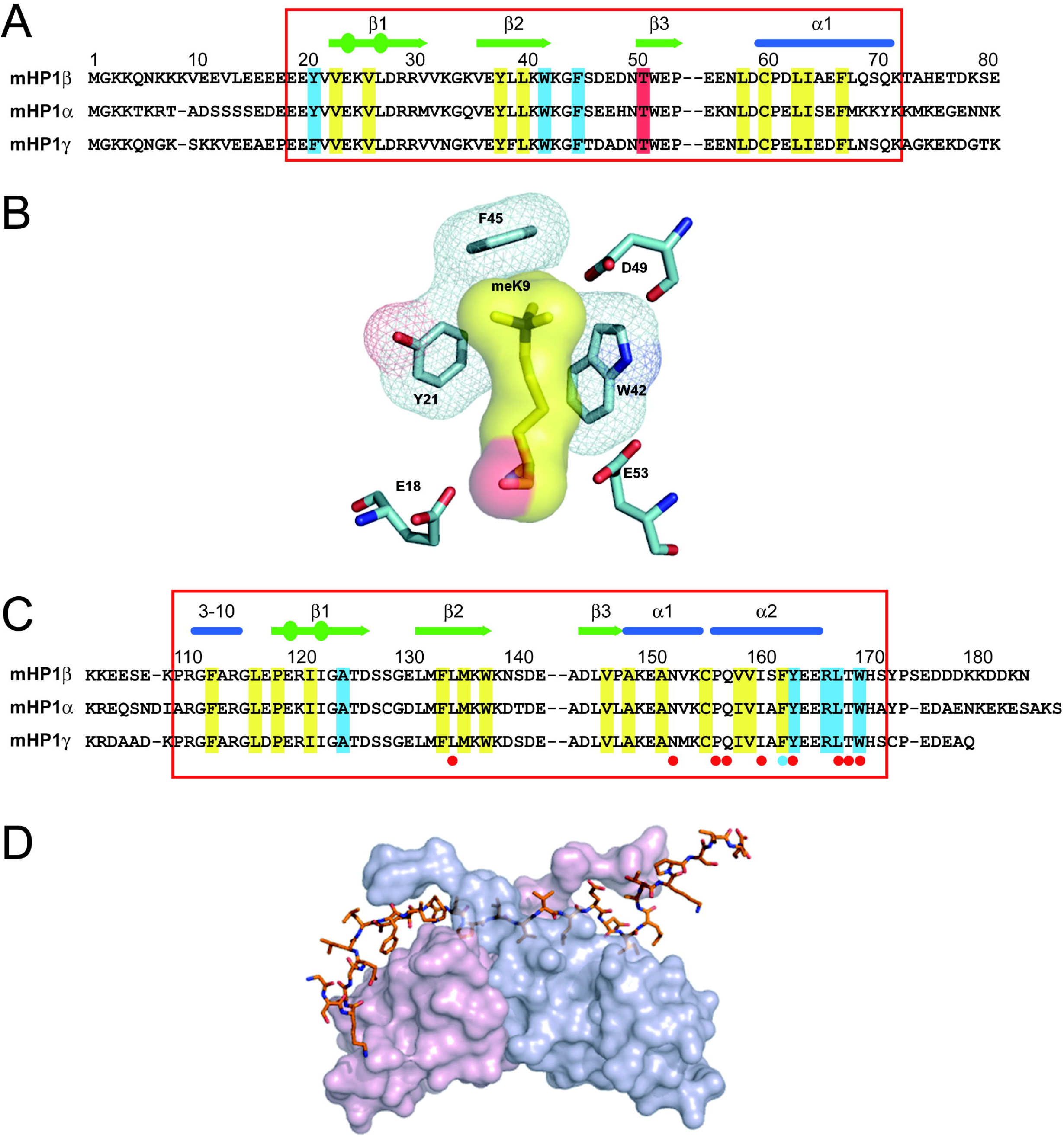
Alignment of peptide sequences of the murine HP1 CDs (A) and the CSDs (C), structure of the aromatic “cage” formed around the K9me3 moiety (B) and the CSD dimer bound to PxVxL motif in CAF-1 peptide (D). **(A)** The CDs from top to bottom are HP1β (aminoacids 1-80), HP1α (amino-acids 1-80) and HP1γ (amino-acids 1-80). The red box denotes the structured part of the CD. The secondary structure elements of the HP1β CD are displayed above the sequence: blue cylinders represent a α-helix (α1) and green arrows represent β-strands; circles within the arrows indicate β-bulges. The residues that make up the hydrophobic core of the CD are shaded in yellow and the aromatic residues that form a notional “cage” around the methyl lysine are given in blue. The Thr51 residue that is phosphorylated after DNA damage [70] is shaded in red. **(B)** Binding of the HP1β CD to H3K9me3 a notional aromatic ‘cage’ is formed from three conserved aromatic residues: Tyr21, Trp42 and Phe45. The interaction between the methylammonium moiety and the aromatic cage is largely electrostatic where the positively charged (cation) moiety is attracted to the negative electrostatic potential of the aromatic groups’ π-system [36]. **(C)** The CSDs from top to bottom are HP1β (amino-acids 103-185), HP1α (amino-acids 106-191) and HP1γ (amino-acids 97-173). The red box denotes the structured part of the CSD. The secondary structure elements of the HP1β CSD are given above the sequence, where the cylinders represent the α-helices (α1 and α2) and the arrows represent β-strands; circles within the arrows indicate β-bulges. The residues that make up the hydrophobic core of the CSD are shaded in yellow and show good alignment with those found in the CD indicating a similar overall structure of the CSD to the CD. Residues that are involved in binding the PxVxL motif are shaded in blue; there is a blue dot below the Phe163 residue, which is involved both in the structure of the CSD and in binding to the peptide. There are red dots below the residues that are involved in CSD:CSD dimerisation. **(D)** Surface view of the CSD homodimer (one monomer in pink and the other in blue) bound to the CAF-1 peptide (shown as a stick model) containing the PxVxL motif, which is involved in intermolecular β pairing with both monomers [37]. Taken and modified from [235].

The interaction of HP1 proteins with H3K9me3 has been resolved at the atomic level and can be illustrated using mammalian HP1β, an archetypal HP1 protein [29]. The primary structure of HP1β is identical in human and mouse [30]. It is essential; HP1β null mutant mice die at birth [31]. HP1β is small at around 25kD having an N-terminal CD and a sequence-related domain towards the C-terminus called the chromo shadow domain (CSD) [32] (*c.f*. Figures 1A vs 1C). These two domains are likely to have arisen by gene duplication [33] and are separated by a less-well conserved “hinge” region (HR) that is flexible and lacks a defined structure [34]. Both the CD and the CSD represent globular protein modules with a diameter of around 30Å. The CD binds the methylated H3K9 tail [35] where three conserved aromatic residues: Tyr21, Trp42 and Phe45 form an “aromatic cage” around the methyl-ammonium moiety (Figure 1B). Most of the binding energy is driven by cation–π interactions where the cation methyl ammonium moiety is attracted to the negative electrostatic potential of the aromatic groups’ π-system [36]. The HP1β CSD dimerizes in solution with the dimer centering upon helix α2 (Figure 1D), which interacts symmetrically and at an angle of 35° with helix α2 of the adjacent CSD subunit and forms a non-polar pit that can accommodate penta-peptides with the consensus sequence motif PxVxL that is found in HP1-interacting proteins [37,38] (Figures 1C and D).

HP1 and H3K9me3 co-localise *outside* constitutive heterochromatin as constituents of heterochromatin-*like domains* and *complexes* that are thought to regulate chromatin-templated processes [30,39,40]. There is evidence that domains/complexes are widespread in mammalian genomes [41] but the question of their number, size and function remains open. Here, we address this question and extend our answer using bioinformatics approach that interrogates the genomes of fission yeast, fruit fly, mouse and human. We focus on domains/complexes that are likely to play critical roles during mammalian development and discuss how segregation of micro-phase separated domains/complexes could drive the compartmentalisation observed in Hi-C experiments. Treating domain/complexes as blocks in a block copolymer (BCP) that micro-phase separate from euchromatin implies that domains/complexes possess a value for χ, the Flory-Huggins parameter. We present a simple theoretical “clutch” model, where the “*unit of incompatibility*” of chromatin is an oligo-nucleosomal “clutch” of 2 to 10 nucleosomes, which could provide an approach for determining χ for domains/complexes experimentally. The magnitude of the product of χ and the number of clutches (N) in a domain/complex (χN) specifies the degree of phase separation of the domain/complex from euchromatin.

## 2. Heterochromatin-*like* domains /complexes in eukaryotes

As a first step towards determining the number and size of heterochromatin-*like* domains/complexes we first investigated the degree to which HP1 and H3K9me3 co-localise *outside* constitutive heterochromatin. This was done for four (distantly-related) genomes, namely man, mouse, *Drosophila melanogaster* and fission yeast. When constitutive heterochromatin is excluded, Pearson correlation coefficients for the co-localisation of HP1α, β and γ with H3K9me3 across the rest of the human genome are 0.73 (α; human H1 ES cells), 0.74 (β; 293T cells) and 0.77 (γ; human H1 ES cells) (Figures 2A-C). The same analysis in mouse ES cells gave correlations of 0.63 for HP1α, 0.69 for HP1β and 0.71 for HP1γ (Figures 2D-F). The mouse results are in agreement with a previous study, which showed that HP1β is preferentially targeted to genomic regions with high local concentrations of H3K9me3 in murine ES cells (correlation coefficient 0.77; [29]). In *Drosophila melanogaster* ovaries the correlation of HP1a with H3K9me3 outside constitutive heterochromatin has a coefficient of 0.92 (Figure 2G). In fission yeast the correlation of Swi6^HP1^ with H3K9me3 is weaker at 0.53 (Figure 2H). This likely reflects the finding that only a limited set of loci outside constitutive heterochromatin are marked by both H3K9me3 and Swi6^HP1^, including the mating-type region, a variety of repeat elements and a number of meiotic genes [18]. If the correlation is made over the entire fission yeast genome the correlation coefficient increases to 0.93 (Figure 2I). For constitutive heterochromatin alone the coefficient approaches unity (0.99; Figure 2J).

**Figure 2:**
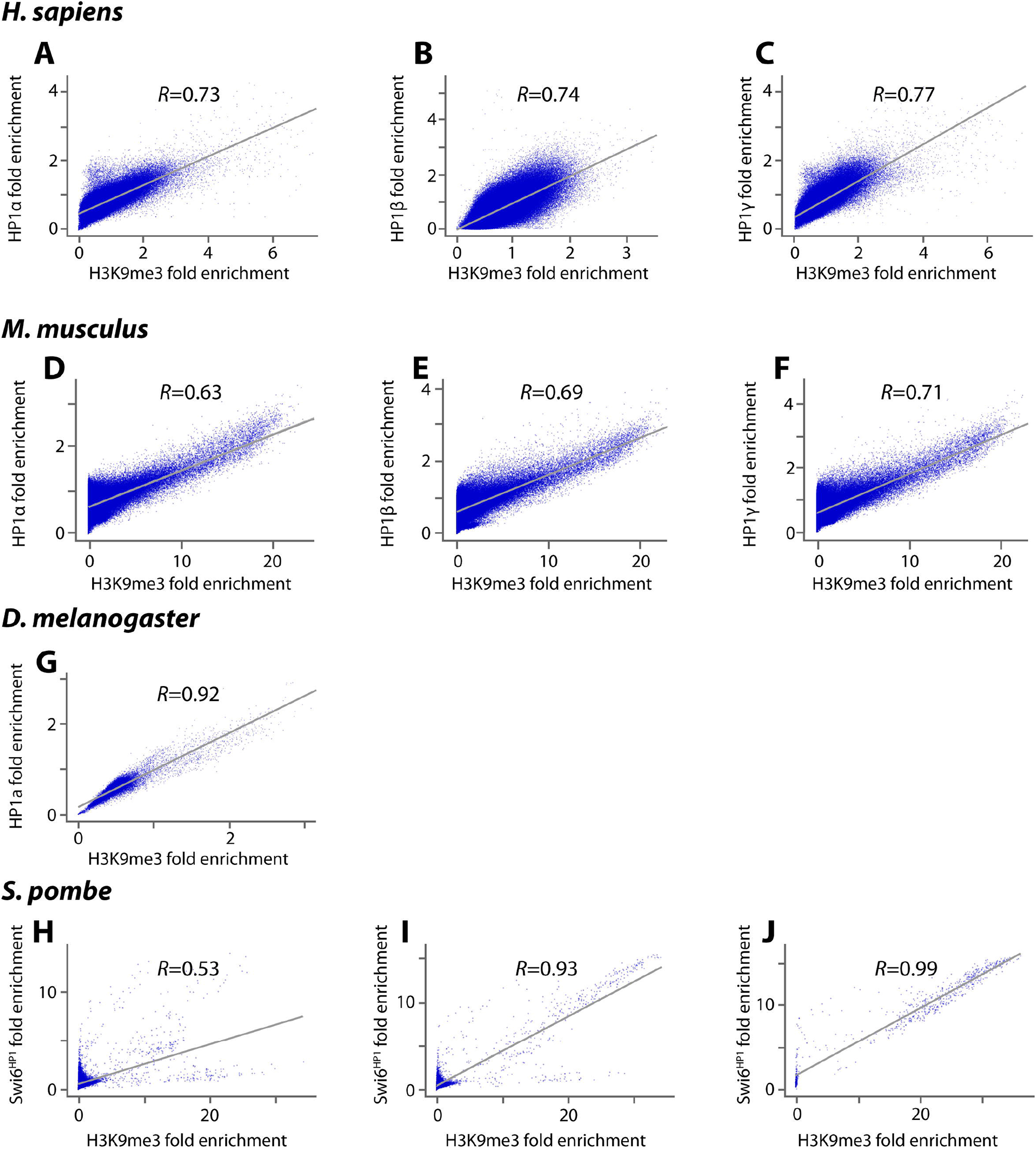
Plots of genome-wide correlation of HP1 proteins and H3K9me3 outside constitutive heterochromatin. The figure depicts fold enrichment of ChIP-seq genome profiles correlated using a 5 kb-sized window for *H.sapiens, M. musculus, D. melanogaster* and a 200 bp-sized window for *S. pombe*. Unless otherwise stated constitutive heterochromatin regions were excluded prior to analysis. **A - C** show the correlations of HP1α (H1 ES cells), HP1β (293T cells) and HP1γ (H1 ES cells) with H3K9me3 in human cells. **D – F** show correlations of HP1α, HP1β and HP1γ with H3K9me3 in mouse ES cells; **G** shows correlation of *Drosophila* HP1a with H3K9me3 in ovaries; **H** shows correlation of Swi6^HP1^ with H3K9me3 in *S. pombe*; **I** shows correlation of HP1β with H3K9me3 over the whole genome of *S. pombe*; **J** shows correlation of Swi6^HP1^ with H3K9me3 over heterochromatin of *S. pombe*.

These data indicate there is a correlation of HP1s with H3K9me3 outside constitutive heterochromatin but that correlation is not absolute. This is consistent with the observation that HP1 does not always “follow code” and can localise to (hetero)chromatin in the absence of H3K9me3 [42]. For example, in the mouse, the ChAHP complex that contains HP1 represses gene expression locally by establishing inaccessible chromatin around its DNA-binding sites and does not depend on H3K9me3-modified nucleosomes [43]. Similarly, in *Drosophila*, a variety of protein partners can localise HP1a to euchromatic sites in the absence of H3K9me3 [44,45]; localisation of HP1a to telomeric constitutive heterochromatin in *Drosophila* is independent of the presence of H3K9me3 at the telomere [21,46]. Alternative modes of HP1 binding to chromatin that do not involve H3K9me3 have also been documented, for example, the HP1 CD or CSD with the H3 histone ‘core’ [47–49], binding of the HR region to DNA and RNA [50–52] and a non-specific electrostatic interaction of the HP1 N-terminal extension with the H3 tail [29]. Of note is the recent demonstration that the Swi6^HP1^ CSD dimer binds to the H2Bα1 helix where it is thought to destabilise the nucleosome and promote phase separation of constitutive heterochromatin in fission yeast [53]. Heterochromatin-*like* domains/complexes lie *outside* constitutive heterochromatin and our survey (Figures 2A-J) indicates that the hallmarks of constitutive heterochromatin (HP1 and H3K9me3) co-localise at many sites where such domain/complexes are assembled. HP1 proteins bound to H3K9me3-marked domains/complexes are constantly exchanging with unbound HP1 proteins in the nucleoplasm. Almost the entire pool of HP1 proteins outside constitutive heterochromatin turns over in around 10 seconds (t_1/2_ = 1-10 seconds; [54,55]); constant exchange maintains compaction of domains/complexes.

We next determined the number and size of the domains/complexes in the four organisms used for the correlation analysis where domains/complexes were put into three categories according to size. Domains are >1Mb and between 0.1 to 1Mb with complexes less than 0.1Mb down to 10kb (Table 1). Our survey showed that the number and size of heterochromatin-*like* domains/complexes increases from fission yeast to fruit flies with a sharp increase from insects to mammals. For human cells we used two different cell lines to estimate the number of HP1α/β/γ-containing domains/complexes. For the human H1 ES cell the HP1α and γ distributions were intersected with H3K9me3 for the whole genome and the genome outside constitutive heterochromatin (Table 1). This showed that outside constitutive heterochromatin there were 163 HP1α/γ-containing heterochromatin-*like* domains and 18853 HP1α/γ-containing complexes, with 90% of the complexes being in the range 10-30kb (Table S1). In human 293T cells there were 859 HP1α/β-containing domains and 32292 HP1α/β-containing complexes, with 75% of the complexes in the range 10-30kb (Table S1). We conclude there are, conservatively, between 163-859 heterochromatin-*like* domains and 18853-32292 complexes in humans depending on cell type. These values may be an underestimate because we could only mine data for two HP1 isoforms per human cell line. For the mouse, we obtained data for all three HP1 isoforms in a single ES cell line, which revealed no domains greater than 1Mb but outside heterochromatin there were 622 HP1α/β/γ-containing heterochromatin-*like* domains between 0.1 to 1Mb and 10227 HP1α/β/γ-containing heterochromatin-*like* complexes, with around 60% of the complexes being in the range 10-30kb (Table S1). In *Drosophila* there are 2 heterochromatin-*like* domains outside heterochromatin in addition to 161 complexes. In fission yeast a survey of the whole genome reveals only 1 heterochromatin-*like* domain that is larger than 0.1Mb that most probably represents the centromeric constitutive heterochromatin of cen3 that is 110 kb in size [56]. Outside constitutive heterochromatin there are only 20 complexes of which the mating type region would be one [18].

**Table 1:** Heterochromatin-*like* domains/complexes in *Homo sapiens, Mus musculus, Drosophila melanogaster* and *Schizosaccharomyces pombe*. In man, the size and number of domains (>1Mb and between 0.1-1Mb) and complexes (0.01-0.1Mb) were determined using ChiP-seq data for H1 ES cells (HP1α+ HP1γ+H3K9me3) [214,243] and 293T cells (HP1α+HP1β+H3K9me3) [244,245]. For the mouse, the same analysis was undertaken in a single cell line, namely mouse ES cells (HP1α+HP1β+HP1γ+H3K9me3) [43]. Data from ovaries were used for the fly (HP1a+H3K9me3) [246]. A culture of fission yeast cells was used to generate the ChIP-seq (Swi6^HP1^+H3K9me3) [247] that were mined for our survey. Domains and complexes were calculated for the whole genome (including constitutive heterochromatin) and the genome without constitutive heterochromatin.

| Organism | Size, Mb | Cell type \| Components | Whole genome | Heterochromatic regions excluded |
| --- | --- | --- | --- | --- |
| *H.sapiens* | >1 | H1 ES cells<br>(HP1α+ HP1γ+H3K9me3) | 48 | 4 |
| | 0.1-1 | H1 ES cells<br>(HP1α+ HP1γ+H3K9me3) | 345 | 159 |
| | 0.01-0.1 | H1 ES cells<br>(HP1α+ HP1γ+H3K9me3) | 19550 | 18853 |
| | >1 | 293T cells<br>(HP1α+HP1β+H3K9me3) | 24 | 4 |
| | 0.1-1 | 293T cells<br>(HP1α+HP1β+H3K9me3) | 1027 | 855 |
| | 0.01-0.1 | 293T cells<br>(HP1α+HP1β+H3K9me3) | 33754 | 32292 |
| *M. musculus* | >1 | ES cells<br>(HP1α+HP1β+HP1γ+H3K9me3) | 0 | 0 |
| | 0.1-1 | ES cells<br>(HP1α+HP1β+HP1γ+H3K9me3) | 1059 | 622 |
| | 0.01-0.1 | ES cells<br>(HP1α+HP1β+HP1γ+H3K9me3) | 12675 | 10227 |
| *D. melanogaster* | >1 | Ovaries (HP1a+H3K9me3) | 7 | 0 |
|  | 0.1-1 | Ovaries (HP1a+H3K9me3) | 27 | 2 |
|  | 0.01-0.1 | Ovaries (HP1a+H3K9me3) | 183 | 161 |
| *S.pombe* | >1 | (Swi6<sup>HP1</sup>+H3K9me3) | 0 | 0 |
|  | 0.1-1 | (Swi6<sup>HP1</sup>+H3K9me3) | 1 | 0 |
|  | 0.01-0.1 | (Swi6<sup>HP1</sup>+H3K9me3) | 23 | 20 |

Our survey indicates that heterochromatin-*like* domains/complexes are widespread in the genomes of (distantly related) eukaryotes (Table 1). At the outset they were proposed primarily as a *general* mechanism for regulating chromatin-templated processes outside constitutive heterochromatin and secondarily as a mechanism for a *special* case where chromosomes and genes exhibit allele-specific parent-of-origin-specific behaviour [30,39,40,57]. Initial support for the general case came from immunofluorescence studies [58] and soon thereafter from the discovery that the universal co-repressor of KRAB-zinc finger proteins (KRAB-ZFPs), KAP1, recruits HP1 proteins to form localised heterochromatin-*like* domains/complexes [59,40]. At around the same time it was shown that the function of HP1-containing heterochromatin-*like* domain/complexes was conserved from fission yeast to man (Figure 8 in [16]). Detailed studies on the KRAB-Zinc finger gene (KRAB-ZNF) clusters on human chromosome 19 have provided insight into how large heterochromatin-*like* domains (up to 4Mb; [60,61]) can be nucleated at particular sites by small heterochromatin-*like* complexes (~6kb; [41]). The domains so assembled make far *cis*-contacts to generate the B4 sub-compartment detected in Hi-C maps [41,62]. An intriguing characteristic of the heterochromatin-*like* domains that encompass the KRAB-ZNF clusters is that the genes within the clusters remain expressible [63]. The domains are thought to ‘protect’ the KRAB-ZNF gene repeats as they have expanded during evolution by preventing illegitimate recombination [60], rather than to repress and silence the repeats. It would seem that heterochromatin-*like* domains/complexes have a variety of chromatin-templated functions an observation supported by studies on HP1, which have shown it to be associated with gene activation as well as with loci involved in other nuclear functions, including transcriptional elongation, RNA splicing and DNA repair [64–71].

Support for the special case alluded to above came from the observation, to be described, that KRAB-ZFPs assemble heterochromatin-*like* complexes at “imprinted genes” that exhibit parent-of-origin-specific gene expression [72].

## 3. Heterochromatin-*like* complexes and preservation of DNA methylation at imprinted gDMRs during pre-implantation embryogenesis

Evidence that phenotypic traits could be subject to parent-of-origin effects came not long after the re-discovery of Mendel’s laws of inheritance in 1900 [73], although Mendel himself thought it indubitable that reciprocal crosses were equivalent, saying: “*…it is perfectly immaterial whether the dominant character belongs to the seed-bearer or to the pollen parent; the form of the hybrid remains identical in both cases*” [74]. We now know of many instances where this is not the case, where the behaviour of chromosomes and genes are dependent upon ancestry. Parent-of-origin-specific behaviour of chromosomes was first observed in insects, in *Sciara* (reviewed by Metz in 1938 [75]) and *Coccidae* [76]. In mammals, pronuclear transfer experiments confirmed that the parental contributions to the zygote were genetically but not functionally equivalent [77,78]. These experiments led to the suggestion that the expression of certain genes, called imprinted genes, was dependent upon parental origin. There are now known to be around 100 imprinted genes that exhibit mono-allelic parent-of-origin-specific expression in mouse and man [79,80]. Such genes are said to be subject to genomic imprinting [81,82]. Genomic imprinting results in genes (or gene clusters) that are either maternally- or paternally-imprinted. Maternally-imprinted genes are associated with a maternal-specific “mark” that acts *in cis* such that there is a heritable (cell-to-cell) change in the behaviour of the gene. The same is true for paternally-imprinted genes except that the “mark” is specific for the paternal allele. Genomic imprinting is necessarily reversible, thus epigenetic, because the parental alleles of an imprinted gene are marked differently in the soma, but the marks must be erased in the germ-line so that both alleles can then be marked again, this time according to the sex of the parent. Imprinting (“marking”) takes place when the parental genomes are separate, which occurs in the respective germ-lines and during the brief period when the pronuclei lie separately in the ooplasm of the newly-fertilized zygote.

In the mouse, depletion of DNA methylation of cytosine (5mC) in CpG dinucleotides leads to dysregulation of genomic imprinting [83]. Since differences in 5mC could be traced back to the sperm and egg it was concluded that the parent-of-origin-specific “mark” is DNA methylation [84]. Parent-of-origin-specific differences in CpG methylation are called gametic or germline differentially methylated regions (gDMRs) and fall within a broader category of CpG-rich genomic regions called CpG islands (CGIs; [85]) that are widespread in the genome, with ~70% of annotated gene promoters in man being associated with a CGI [86]; within the bounds of definition both maternally- and paternally-imprinted gDMRs are recognisable as CGIs [87,88]. CpGs that are part of CGIs are usually unmethylated, whether the associated gene is active or inactive [85], but imprinted gDMRs are exceptions where CGIs are methylated. Accordingly, classical imprinted gDMRs are methylated in either the female (maternally-imprinted) or the male (paternally-imprinted) germline and retain this parent-of-origin specific methylation following fertilization and during pre-implantation development [82]. In the mouse, there are ~26 (23 maternal and 3 paternal) definitive imprinted gDMRs [89,90] (Figure 3A).

**Figure 3:**
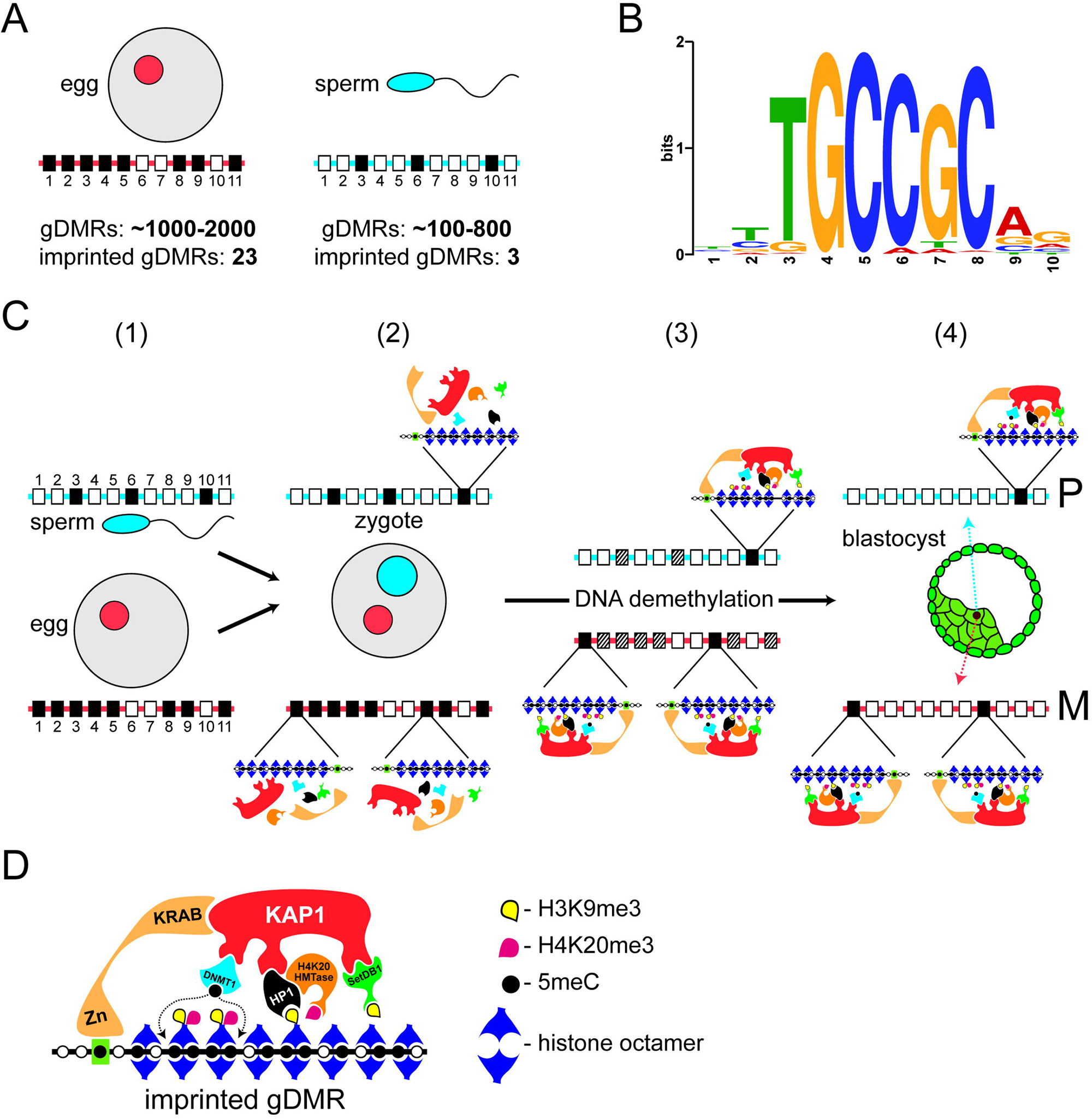
Preservation of DNA methylation at imprinted gDMRs by a localized heterochromatin-*like* complex. **(A). There are many more (imprinted and non-imprinted) gDMRs than definitive imprinted gDMRs.** The oocyte nucleus (red circle) contains 1-2000 oocyte-specific gDMRs of which 23 are definitive imprinted gDMRs. The sperm nucleus (blue circle) contains around 100–800 gDMRs of which 3 are definitive imprinted gDMRs. Below the oocyte and sperm are schematic maternal and paternal homologous chromosomes that carry CpG islands (CGIs) depicted as rectangles numbered 1 through to 11 on red (maternal homologue) and blue (paternal homologue) lines. Open rectangles represent non-methylated CGIs. Some methylated CGIs are shared (*e.g*. closed rectangle at position 3 on both parental homologues) and are not gDMRs. Some methylated CGIs are non-imprinted gDMRs (closed rectangles at position 6 on paternal chromosome and positions 2, 4, 5, 9 and 11 on the maternal chromosome) that will lose their methylation during the DNA demethylation that takes place as embryos pass through preimplantation development. A few methylated CGIs are imprinted gDMRs (closed rectangles at position 10 on the paternal chromosome and 1 and 8 on the maternal homologue) that will retain their methylation status through DNA demethylation. As explained, there are around 1-2000 non-imprinted and imprinted gDMRs present in the oocyte nucleus and around 100-800 non-imprinted and imprinted gDMRs will enter with the sperm (see text for detailed numbers). This difference in number in respective germ cells is reflected in the difference in closed rectangles on the maternal (red line) and paternal (blue line) homologues. **(B). Consensus binding site of ZFP57.** The TGCCGC hexamer motif is found in imprinted gDMRs and bound by ZFP57 when the central CpG dinucleotide is methylated. This motif shown (based on [117]) was downloaded from the Tomtom database (http://meme-suite.org/tools/tomtom) and has the searchable identifier ZFP57_MOUSE.H11MO.0.B. **(C) Preservation of methylation at imprinted gDMRs. (1)** The paternal (sperm nucleus in blue) and maternal (oocyte nucleus in red) nuclei contain homologous chromosomes that carry CpG islands (CGIs) depicted as described in **(A)**. (**2)** The maternal (in red) and paternal (in blue) pro-nuclei contain the homologous chromosomes (red and blue lines respectively) described in **(1)**. Of the ~1600 non-imprinted and imprinted gDMRs in the zygote, only a small percentage - the imprinted gDMRs – will preserve DNA methylation in the face of the DNA demethylation that takes place during pre-implantation development [98]. The initial assembly of the heterochromatin-like complexes that preserve methylation at imprinted gDMRs takes place in the newly-fertilized zygote (see text for details). It should be noted that H4K20HMTases have only been detected in the zygote by RT-PCR owing to lack of specific antibodies so it remains to be shown that the mRNAs are translated to give active proteins. (**3)** Preservation of methylation at imprinted gDMRs on the paternal (position 10) and maternal (positions 1 and 8) homologues is due to localized heterochromatin-like complexes at imprinted gDMRs. We have included H4K20HMtases as part of the heterochromatin-like complex that generate H4K20me3 at the imprinted gDMRs because trace amounts have been detected by RT-PCR albeit translated protein has yet to be shown. The complexes preserve DNA methylation at imprinted gDMRs throughout pre-implantation development; non-imprinted gDMRs and methylated CGIs become de-methylated (stippled rectangles). **(4)** Global levels of DNA methylation reach their lowest point in embryonic nuclei of the blastocyst. However, methylation at imprinted gDMRs is preserved by the heterochromatin-like complexes shown in **(3)**, on the paternal (position 10) and maternal (position 1 and 8) homologues. P denotes paternal homologue and M the maternal homologue. Taken and modified from [236]. **(D) Assembly of localized heterochromatin-like complex at imprinted gDMRs.** Methylation of cytosines in CpG dinucleotides (black circles) is preserved by the assembly of a heterochromatin-like complex at imprinted gDMRs. The complex is targeted by the KRAB zinc-finger protein ZFP57 that binds the hexamer motif TGCCGC when the cytosine in the CpG is methylated (black circle in green rectangle). This in turn recruits KAP1, which is a modular protein that acts as a focal point for the recruitment of Setdb1 histone methyltransferase, HP1 and Dnmt1. HP1 binds the H3K9me3 generated by Setdb1 and recruits a H4K20me3 histone methyl-transferase that generates H4K20me3 thus forming the H3K9me3:HP1:H4K20me3 pathway. DNA methylation at the imprinted gDMR is maintained (dotted lines) by Dnmt1. Taken and modified from [236].

The identification of a small number of imprinted gDMRs during the pre-genomic era prompted efforts directed towards identifying hypothetical specialised sequence elements that would be recognised by specific *trans*-acting factors that target the *de novo* DNA methylation machinery to imprinted gDMRs during gametogenesis. However, later whole genome studies revealed a very different picture, which led to the abandonment of the notion that specific imprinting machinery operates in the germline. Mining the sperm and oocyte methylomes showed that there were many more gDMRs in the gametes, far above the number of definitive imprinted gDMRs. Depending on the study, the mouse oocyte nucleus contains around 1-2000 oocyte-specific gDMRs, with the sperm-specific gDMRs numbering between 185 and 818 [91,92] (Figure 2A). The combined total is roughly1600 (imprinted and non-imprinted) gDMRs in the pro-nuclei of the newly fertilised zygote [92]. This is almost two orders of magnitude greater than the number (~26) of classical definitive imprinted gDMRs whose methylation is retained after fertilisation and through pre-implantation embryogenesis [89,90] (Figure 2A). Putting it short, imprinted gDMRs are not uniquely targeted for DNA methylation; imprinted and non-imprinted gDMRs are methylated by mechanisms common to CGIs [93,94]. The difference is that allele-specific methylation at imprinted gDMRs is preserved after fertilisation while the differential methylation at non-imprinted gDMRs is not.

Pre-implantation embryogenesis in mouse and man is characterised by a global DNA demethylation of the parental genomes [95,96]. Demethylation is thought to ensure that the embryonic epigenome is purged of any barriers to pluripotency, which is essential for those cells of the inner cellular mass that will go on to form the tissues and cell types of the embryo proper [97]. DNA methylation at the imprinted gDMRs is preserved during this DNA demethylation phase through the assembly of localised heterochromatin-*like* complexes at imprinted gDMRs [98,99]. It is the sequence-specific assembly of heterochromatin-*like* complexes at imprinted gDMRs that is the key to understanding genomic imprinting in mammals, consistent with the earlier proposal [30].

Preservation of methylation at imprinted gDMRs requires binding of KRAB zinc-finger proteins (KRAB-ZFPs), ZFP57 [72,100] and ZFP445 [101]. Methylation of all imprinted gDMRs, except one, is lost in ZFP57/445 mouse double mutants [101]. Characterisation of ZFP57 in the mouse has shown that binding of ZFP57 is methylation sensitive with its binding being a hexamer motif TGCCGC found in imprinted gDMRs, where the central CpG dinucleotide is methylated [100,102,103] (Figure 2B). Assembly of ZFP57/445-directed heterochromatin-*like* complexes is likely to take place soon after fertilization since most of the constituents are laid down maternally (Figure 2C). In mouse oocytes there are maternal stores of ZFP57 and KAP1 and their loss affects methylation at imprinted gDMRs [72,104]. The Setdb1 HMTase that interacts with KAP1 is localised to peri-nucleolar rims of the pro-nuclei in the zygote [105] and maternal deletion of Setdb1 leads to a dramatic defects in preimplantation development [106]; deletion in ES cells leads to DNA demethylation of imprinted gDMRs [107]. Setdb1 generates the H3K9me3 binding site for HP1 proteins that are also found in the oocyte cytoplasm and pro-nuclei of the early embryo [108,109]. The maintenance DNA methyltransferase Dnmt1that interacts with KAP1 [100,110] is barely detectable in the ooplasm and in the early embryo [111,112]. Yet these trace amounts are essential for methylation of imprinted gDMRs because loss of maternal-zygotic Dnmt1 leads to loss of methylation at all imprinted gDMRs [113]. H4K20me3 catalysed by H4K20HMTases during oogenesis is present in the maternal pronucleus but is undetectable by immunofluorescence in nuclei at later preimplantation stages [109,114,115]. It has yet to be shown whether the (very) low levels of H4K20HMTase mRNAs present in the zygote and pre-implantation stages [114] are translated into active proteins and generate H4K20me3 at imprinted gDMRs though interactions with the resident heterochromatin-*like* complexes.

Once assembled, the complex preserves methylation at imprinted gDMRs throughout the demethylation phase, which is complete at the blastocyst stage where the lowest levels of global DNA methylation are reached (Figure 3C) [94,95,116]. Biochemical and functional studies using ES cells (derived from the blastocyst) have shown that the structural proteins and enzymatic activities of the heterochromatin-*like* complexes are present at imprinted gDMRs [100,107,117–119] (Figure 3D). H4K20me3 is clearly detected at imprinted gDMRs in ES cells [117], indicating the recruitment of an H4K20HMTase [118], most likely through a known interaction of H4K20HMTases with HP1 [120] (Figure 3D).

There are two ways by which heterochromatin-*like* complexes could preserve DNA methylation at imprinted gDMRs. The first is by protecting imprinted gDMRs from the activity of demethylating Tet dioxygenases that are present in the early embryo [97,98,121]. The second is mediated by the interaction of KAP1 with Dnmt1 [100,110] (Figure 3D), which is important because Dnmt1 is scarce during the DNA demethylation phase of pre-implantation development [111–113]. The KAP1-Dnmt1 interaction would have the effect of concentrating trace amounts of Dnmt1 in the vicinity of imprinted gDMRs thereby ensuring maintenance of 5mC (Figure 3C). Dnmt1 may also be recruited to imprinted gDMRs through a known interaction with HP1 proteins that has been shown to increase local DNA methylation levels [122].

The size of the heterochromatin-*like* complex at the imprinted gDMRs is around 6kb [60,61,119]. The mechanism(s) by which the size of heterochromatin-*like* complexes is regulated is of interest because in other regions of the genome small, localised, heterochromatin-*like* complexes, such as that found at the imprinted gDMRs, act as nucleation sites for the assembly of much larger domains. This is the case for KRAB-ZFP-directed heterochromatin-*like* complexes that nucleate the assembly of KRAB-ZNF heterochromatin-*like* domains that can range up to 4Mb in size [41,60].

After the embryos implant into the maternal endometrium early post-implantation development is characterized by elongation of the primitive streak along the epiblast whereupon gastrulation begins and cells of the embryo undergo finely orchestrated morphogenetic movements to form the three germ layers, ectoderm, mesoderm and endoderm [123]. Of note is that around this time H3K9me3-marked heterochromatin is transiently deployed in germ layer cells [124]. This transient deployment is likely involved in an evolutionary restriction observed during vertebrate development called the phylotypic period or progression, to which we now turn.

## 4. Heterochromatin-*like* domains/complexes and the phylotypic progression during vertebrate development

In his popular work *Anthropogenie* Haeckel [125] published some of the most famous pictures in Biology: a series of comparative drawings showing different animals arising from near identical somite-stage embryos. For more than a century there has been controversy over what weight ought to be placed on the images [126], nevertheless, what they illustrated so graphically was there is a stage in development where an animal most closely resembles other species (Figures 4A and B). Ironically, this notion has become one of the central concepts in evolution and development because a similar embryonic stage can be identified in each phylum and has been termed the phylotypic stage [127]. In vertebrates, the identification of a precise phylotypic stage that is identical in all species has been elusive owing largely to the vagaries of heterochrony [128,129]. Rather, there is thought to be a phylotypic “period” [128] or “progression” [130] that roughly corresponds to organogenesis, where numerous, undifferentiated organ primordia are developing from the three germ layers (Figures 4A and B) [130–132]. Here, we use the term “phylotypic progression” because it most closely describes the character of the molecular mechanisms, to be described, that restrict the amount of evolution allowed during this developmental window.

**Figure 4:**
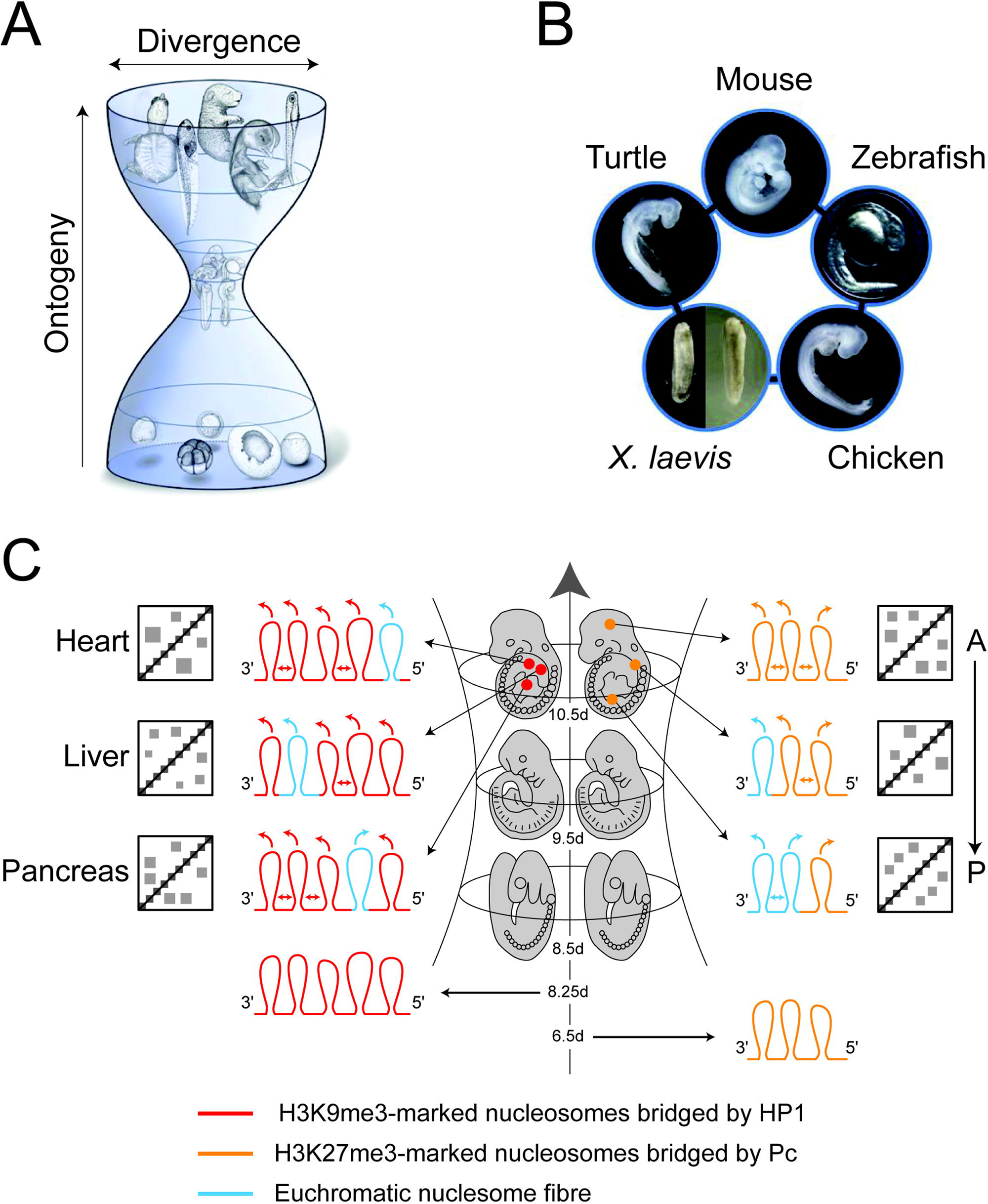
The phylotypic progression of vertebrate development and the generation of cell-type- and position-specific contact enrichments that contribute to epigenetic compartmental domains (ECDs). **(A) The developmental hourglass model for vertebrate development.** The model predicts that the mid-embryonic organogenesis stages (phylotypic progression) represent the developmental stages with highest morphological conservation across vertebrates. The phylotypic progression encompasses the developmental window when the anterior-posterior axis (body plan, *Bauplan*) is laid down. By the end of the phylotypic progression cell-type-specific patterns of gene expression have been initiated and position-specific (*Hox* code) patterns of gene expression are established and thereafter maintained for the rest of development. Taken and modified from [132]. **(B) The phylotypic progression for vertebrates.** This figure is taken from [132] where two stages of *X. laevis* were shown because there was no statistically significant difference between these two stages. **(C) Establishment of cell-type- and position-specific contact enrichments that contribute to ECDs.** Depicted in the middle is the “bottle-neck” of the hourglass depicted in **(A)**, which shows the embryonic stages of the murine phylotypic progression where embryos show greatest similarity with other vertebrates. On the left of the progression is depicted the establishment of cell-type-specific contact enrichments. By day 8.25 of embryonic development H3K9me3-marked heterchromatin is transiently deployed to compact and silence genes that regulate cell-type-specific differentiation. This is shown as five chromatin loops (red loops) that represent H3K9me3-marked chromosome fibers that are “bridged” by HP1 proteins. As embryos pass through the phylotypic progression there is a progressive loss in H3K9me3-marked heterochromatin. By the end of the phylotypic progression there is cell-type-specific loss of H3K9me3-marked heterochromatin and differentiation-specific genes are expressed. Far *cis*- and *trans*-contact between micro-phase separated heterochromatin-*like* domains/complexes result in cell type-specific contact enrichments that emerge as cell-type ECDs (double-headed arrows denote (far) *cis*-interactions; arrows on top of loops indicate *trans*-interactions; the cartoon Hi-C maps show, crudely, cell-type specific contact enrichments that become part of ECDs; A type compartments are not shown). The scenario is similar to the generation of position-specific contact enrichments that become part of ECDs that are shown on the right. On day 6.5 the *Hox* genes clusters are assembled as Pc-G domains (orange loops on the bottom right) that represent a “closed” chromatin domains whose constituents are H3K27me3-marked nucleosomes and the PRC1- and PRC2-complexes, where the Pc homologue in PRC1 “bridges” H3K27me3-marked nucleosomes (Diagram A in Box 1, bottom row). As embryos progress through the phylotypic progression there is a progressive 3’ to 5’activation of *Hox* genes along the *Hox* gene cluster (temporal collinearity). By the time embryos leave the phylotypic progression the spatially-restricted patterns of Hox gene expression (spatial collinearity) have been established so that the *Hox* code for each region of the embryo are stable for the rest of development. In nuclei from the posterior trunk, much of the *Hox* gene cluster is in a euchromatic conformation that facilitates *Hox* gene expression (two blue “euchromatic” loops) with only a small region of the Hox cluster compacted into the remaining Pc-G domain (orange Pc-G loop). The *cis*- (denoted by double-headed arrows) and *trans*- (given by arrows on top of the loops) interaction that are mediated by this configuration of loops contribute position-specific contact enrichments to ECDs that are shown in the Hi-C maps (on the far right of Figure 4C). In nuclei from the mid-trunk region a smaller region of the *Hox* cluster is in a euchromatic conformation (one blue “euchromatic” loops) while a larger region remains compacted into a Pc-G domain (two orange Pc-G loops). This configuration of loops gives rise to position-specific contact enrichments that reflect the position of the cell along the A-P axis. These contact enrichments are shown in the middle Hi-C map cartoon to the right of the loops in figure 4C. In the forebrain where *Hox* genes are not expressed the entire *HoxD* cluster is assembled into a Pc-G domain (three orange loops at top on right in Figure 4C). This configuration of loops give rise to contact enrichments that are specific for this anterior position along the A-P axis and are depicted in the cartoon Hi-C maps at the top right of Figure 4C. The position-specific contact enrichments are a small fraction of the contacts that contribute to ECDs. Not shown in the cartoon Hi-C maps are the myriad of additional contacts between ~2000 Pc-G domains/complexes that are found elsewhere in mouse and human genomes nor are the contacts mediated by heterochromatin-*like* domains/complexes.

The reduced inter-species variability of the phylotypic progression is flanked on either side by an earlier stage of ontogeny at which species differ markedly from one another, and a later stage that shows a progressive divergence among species (Figures 4A and B). The divergence of morphologies on either side of the phylotypic progression formed the basis of the hourglass model of vertebrate development [130–132]. Molecular explanations for the evolutionary “bottleneck” through which embryos pass are of two kinds. One is that there are signalling pathways among developmental modules in the mid-embryonic stages that are highly inter-dependent and make this period developmentally constrained, thus leading to evolutionary conservation [131]. The second relates to the mechanism(s) by which the anterior-posterior (A-P) axis (body plan; *Bauplan*) is laid down, specifically, the mechanisms that regulate the temporal and spatial collinearity of *Hox* cluster gene expression [130], where perturbations in the timing and/or extent of *Hox* gene expression are deleterious and this, again, leads to a restriction in the amount of evolution allowed. Support for this second explanation, as well as for the hourglass model in general, has come from cross-species transcriptome comparisons. Transcriptome profiling of mid-embryonic (around gastrula to organogenesis) stages of four vertebrate species (mouse, chicken, *X. laevis* and zebrafish) showed conserved expression profiles during the above stages including conserved expression of *Hox* genes [133]. The relationship between the spatially-restricted *Hox* gene expression patterns, the laying down of the *Bauplan* and the phylotypic progression reached its apotheosis in the remarkable observation that all three are intimately associated across different phyla. This led to the hypothesis that the association is a universal trait in animals and the defining characteristic of the zootype [134].

We suggest that the major cause for the phylotypic progression in vertebrates is the requirement for properly establishing epigenetic compartmental domains (ECDs; Box 1) so that by the end of the progression both position-specific and cell-type-specific cellular identities are safeguarded. It is a general mechanism that accommodates the role played by *Hox* gene expression in determining position-specific identities along the A-P axis [135]. ECDs contribute to ensuring that they remain so. ECDs are defined as contact enrichments seen in Hi-C maps that are generated by segregation of *micro*-phase separated HP1-containing heterochromatin-*like* and Pc-containing Pc-G domains/complexes (Box 1; [41]). A role for heterochromatin-*like* domains/complexes in regulating the phylotypic progression is indicated from two recent studies. The first, a study of the reprogramming of H3K9me3-marked heterochromatin during early mouse development, showed that in recently implantation embryos (around day 6.5) there is a gradual increase in the association of H3K9me3-marked heterochromatin with lineage-incompatible genes [136]. The second showed that levels continue to rise reaching their maximum in germ layer cells on day 8.25 where there is a net increase in association of H3K9me3-marked heterochromatin with genes that regulate differentiation of adult cell types [124]. Repression of differentiation-specific genes is transient. Beyond day 8.25, as embryos traverse the phylotypic period there is a progressive loss of H3K9me3 and chromatin compaction at many sites in the genome and at those sites tissue-specific gene expression with concomitant differentiation begins [124]. By the time embryos exit the phylotypic progression (we give it as around day 10.5 in Figure 4C) there is cell-type-specific loss of H3K9me3-marked heterochromatin, where previously repressed genes take up euchromatic conformations that promote tissue specific gene expression (blue loops on the left in Figure 4C). Nuclei of differentiating cells still have H3K9me3-marked heterochromatin domains/complexes elsewhere in their genomes and “bridging” within and between H3K9me3-marked nucleosome fibers by HP1 contribute to the emergence of cell-type-specific contact enrichments in ECDs (red loops on the left in Figure 4C and associated cartoon Hi-C maps). ECDs safeguard cellular identity (Diagrams A to C in Box 1). Notably, RNAi “knock-down” screens for genes whose depletion destabilise cellular identity, identified genes that encode CAF-1, the SUMO-conjugating enzyme UBE2i, SUMO2, SETDB1, ATRX and DAXX proteins [137,138]. All are involved in either nucleation or replication of heterochromatin-*like* domains/complexes thus providing a link between safeguarding cellular identity and ECDs (Diagram C in Box1; Figure 4C on left; [41]).

### Box 1: Epigenetic compartmental domains (ECDs) in mammals

Contact enrichments detected in Hi-C maps that are generated by segregation of micro-phase separated heterochromatin-*like* and Pc-G domains/complexes are termed epigenetic compartmental domains (ECDs; [41]). Micro-phase separation of heterochromatin-*like* domains/complexes is caused by “bridging” of H3K9me3-marked nucleosomes by HP1 proteins; micro-phase separation of Pc-G domains/complexes is caused by “bridging” of H3K27me3-marked nucleosomes by Polycomb CBX homologues. Like-with-like segregation of micro-phase separated domains/complexes generates contact enrichments that drive the compartmentalisation detected in Hi-C maps. ECDs overlap with B-type compartmental domains and represent the “epigenetic” component of cellular identity and contribute to safeguarding that identity. The evidence and reasoning that supports this definition of ECDs is detailed below, using as our guide three diagrams.

The conclusion drawn from intersecting data from computational correlation, principle component analysis and ChIP-Seq data was that the cell-type specific checkerboard (or plaid) pattern observed in Hi-C maps represented the folding of chromatin into euchromatin (A-type compartments) and heterochromatin (B-type compartments) [237–239]. Importantly, the H3K9me3 and H3K27me3 epigenetic modifications used to define B-type compartments as heterochromatic are the only histone modifications that are truly epigenetic, *i.e*., heritable from one cellular generation to the next [210]. This is because each modification has its own “*write-and-read*” mechanism (Diagram **A**) that enables HP1-containing heterochromatin-*like* and Polycomb-Group (Pc-G; consisting of H3K27me3, PRC1 and PRC2; [140,141]) domains/complexes to be faithfully inherited from one cell generation to the next.

**Diagram A:**
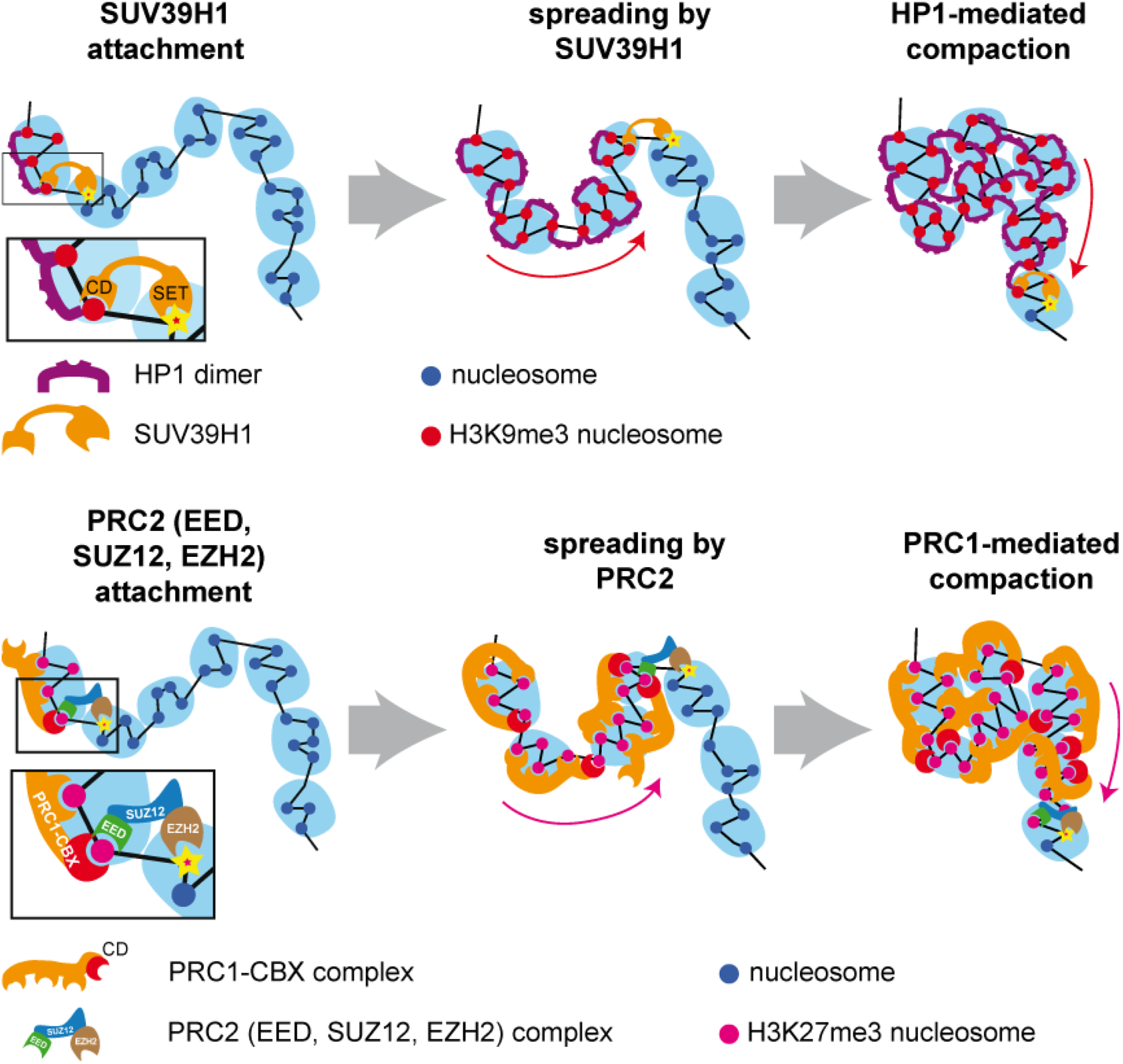
The *write-and-read* mechanisms for the H3K9me3 and H3K27me3 *epigenetic* histone modifications. **Top row:** For the H3K9me3 “write-and-read” mechanism the CD of the SUV39H1 K3K9 HMTase binds H3K9me3 (left), which increases allosteric activation of the SET (methyltransferase) domain that in turn enhances the methylation at H3K9 of neighbouring naïve nucleosomes [240]. The HP1 dimers “bridge” H3K9me3-marked nucleosomes in the wake of the “spreading” H3K9me3 domain, which stabilizes the zig-zag geometry within a “clutch” (middle) with concomitant compaction (right; [29,180]). HP1 bridging of H3K9me3-marked nucleosomes resulting in enrichment (compaction) drives *micro*-phase separation of the heterochromatin-*like* domains/complexes from euchromatin (see Diagram B given next). **Bottom Row:** In the case of H3K27me3, it is the PRC2 complex consisting of the core subunits EED, SUZ12 and EZH2, that regulates the “write- and-read” mechanism. Accordingly, EED binds H3K27me3 and this binding allosterically activates the SET domain of EZH2, which in turn enhances the methylation at H3K27 of neighbouring naïve nucleosomes (left). The CD of a CBX protein that is a homologue of *Drosophila Polycomb* binds in the wake of the “spreading” domain of H3K27me3-marked nucleosomes (middle). The best studied of the mammalian Pc-homologues is CBX2 [187]. CBX2 is a subunit of the PRC1 complex and CBX2 contains a specific domain called the compaction and phase separation (CaPS) domain that compacts (four) nucleosomes [186,187]; the compaction stabilizes the zig-zag geometry within a “clutch” (right). It is the combined action of PRC2-PRC1complexes that drive *micro*-phase separation of Pc-G domains/complexes.

Like-with-like interactions drive segregation of micro-phase separated heterochromatin-*like* domains/complexes (as shown in Diagram **B**); Pc-G domains/complexes would segregate with Pc-G domains/complexes. Segregation involves both *cis*- (shown in Diagram **B**) and *trans*-interactions (not shown in Diagram **B**).

**Diagram B:**
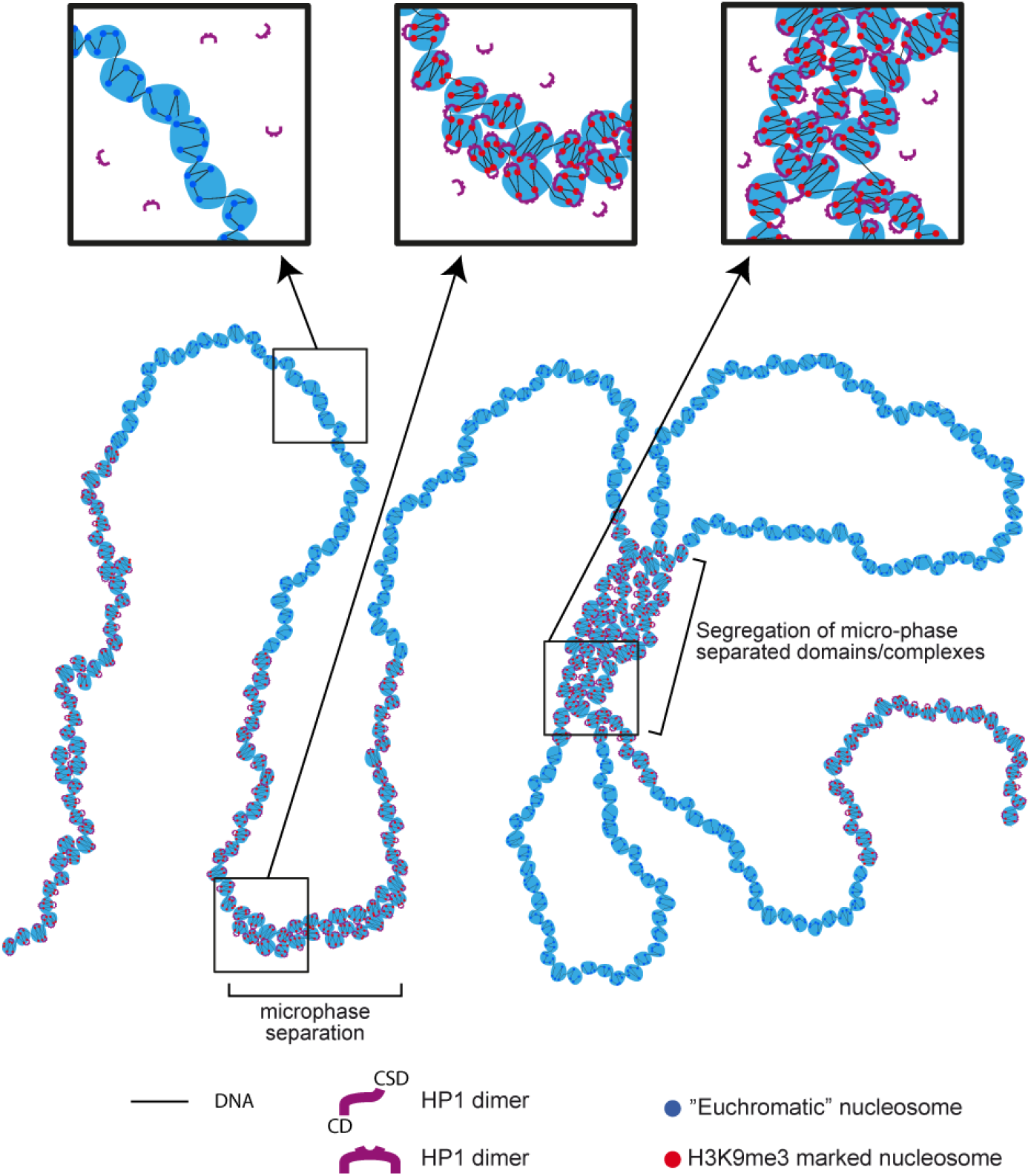
*Micro*-phase separation and segregation of heterochromatin-*like* domain/complexes. Bridging of H3K9me3-marked nucleosomes by HP1 results in *micro*-phase separation of heterochromatin-*like* domains/complexes (middle loop). Bridging results in stabilisation of zig-zag geometry of H3K9me3-marked nucleosomes in heterochromatin-*like* “clutches” (middle box at top of figure). This contrasts with euchromatic “clutches” (box on left) where the nucleosomes are disorganised with only partial zig-zag geometry. Like-with-like attraction (binding potential), due to an entropic effect (see text for details), results in *cis*-(shown in Diagram B) and *trans*- (not shown) interactions between domain/complexes. Interactions are stabilised by bridging of H3K9me3-marked nucleosome fibres by HP1 (box on right) and stabilises segregation of domains/complexes. Contacts resulting from segregation are detected as contact enrichments that contribute to ECDs in Hi-C experiments. Segregation of micro-phase separated domains/complexes is unlikely to be static (as drawn). Micro-phase separated domains/complexes within segregated assemblies will be subject to constant disassociation and association.

C*is*- and *trans*-contacts that result from segregation of heterochromatin-*like* and Pc-G domains/complexes emerge as ECDs in Hi-C maps (Diagram **C**). Notably, screens for genes that are necessary to safeguard cellular identity identified genes that encode CAF-1, the SUMO-conjugating enzyme UBE2i, SUMO2, SETDB1, ATRX and DAXX proteins [137,138]. All are involved in either nucleation or replication of heterochromatin-*like* domains/complexes thus providing a link between safeguarding cellular identity and ECDs [41].

**Diagram C:**
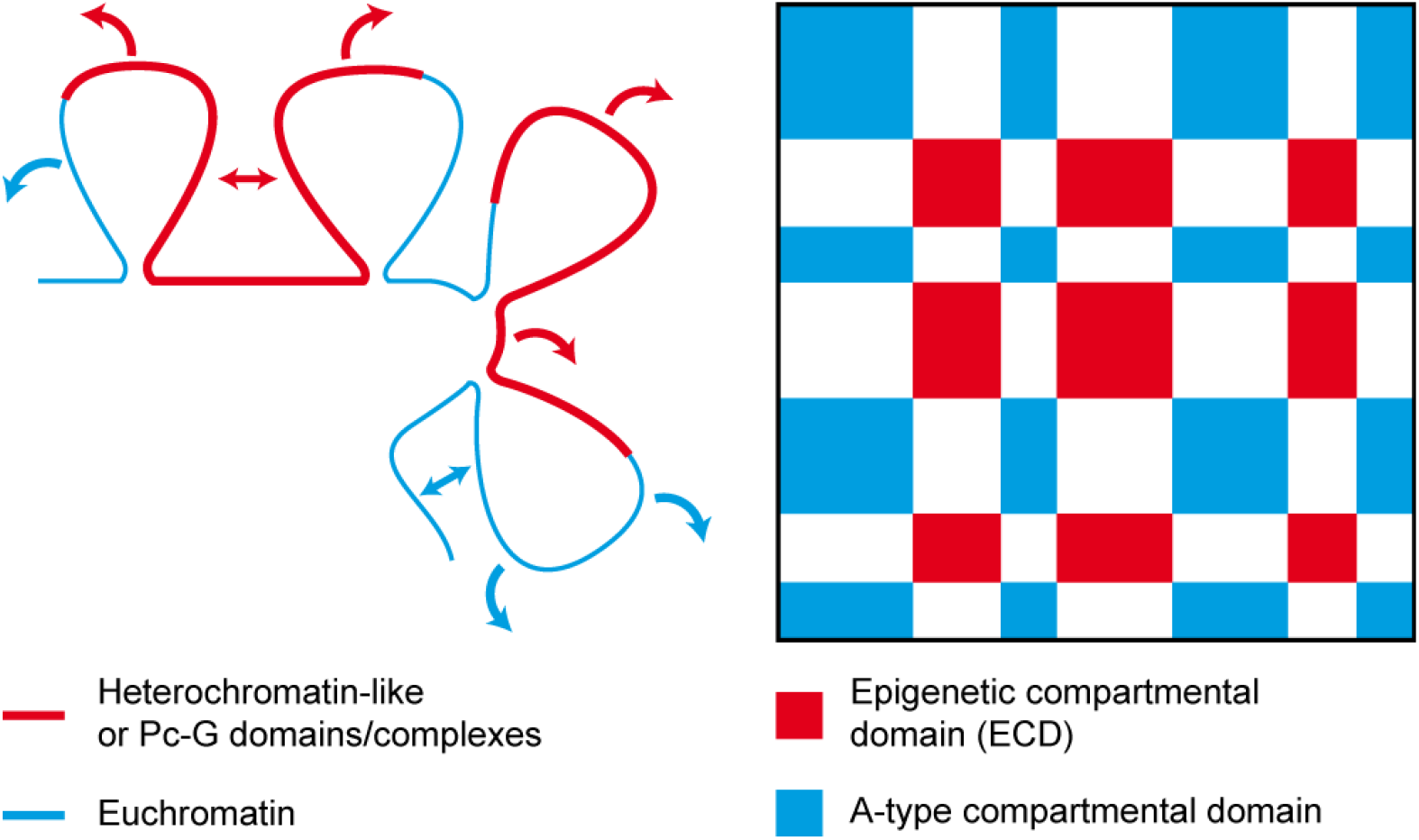
*Cis*- and *trans*- involving micro-phase separated heterochromatinlike or Pc-G domains/complexes generate ECDs. Heterochromatin-*like* <u>or</u> Pc-G domains/complexes (red lines) segregate through like-with-like *cis*- (red double-headed arrows) and *trans*- (red arrows) interactions. These interactions generate contact enrichments that emerge as epigenetic compartmental domains (red EDCs in the cartoon Hi-C map). For euchromatin, *cis*- (blue double-headed arrows) and *trans*- (blue arrows) contacts generate the A-type compartmental domains (blue contact enrichments in cartoon Hi-C map). The cartoon Hi-C map was taken and modified from [239].

B-type compartmental domains are unlikely to be precisely equivalent to ECDs, *i.e*., it is unlikely that B-type compartments are generated solely by contact enrichments that are a consequence of segregation of micro-phase separated heterochromatin-*like* and Pc-G domains/complexes. For example, it is known that B3 sub-compartment includes contact enrichments that result from interactions between lamin-associated loci [182]. HP1 proteins are also known to associate with the nuclear lamina [241] but the degree to which heterochromatin-*like* domains/complexes are involved in interactions between lamin-associated loci that generate these contact enrichments is not known. Functional experiments will be needed to define the precise extent to which ECDs overlap with B-type heterochromatic compartmental domains.

Pc-G domains/complexes also contribute contact enrichments to ECDs (Box 1; Figure 4C on the right) and the best described Pc-G domains are those that compact the *Hox* gene clusters. *Hox* genes are determinants of cellular fate and the positional identities of post-occipital tissues in the mouse (tissues below the skull, including the trunk and limbs) are determined by the collinear expression of *Hox* genes [139]. In cells where *Hox* genes are not expressed, such as ICM-derived ES cells and cells of the post-implantation epiblast, repressive Pc-G domains compact the ~100kb *Hox* gene clusters; constituents of Pc-G domains include the canonical H3K27me3 histone modification and the PRC1 and PRC2 complexes [140,141]. In the forebrain, where *Hox* genes are not expressed, the same H3K27me3-marked domain has been shown to encompass the *HoxD* and *HoxB* clusters [142]. Thus, in cells of the epiblast (around day 6.5 in the mouse), prior to entering the phylotypic progression, the *Hox* gene clusters are assembled into repressive Pc-G domains (depicted on the bottom right of Figure 4C as three orange Pc-G loops).

*Hox* gene expression is observed first on day 7.2 [143] and, once initiated, there is a gradual 3’ to 5’ activation along the *Hox* gene clusters with the 3’-most genes (group 1 genes) of the cluster being expressed first to be followed one after another by *Hox* genes that reside more 5’ finally ending with the 5’-located (group 13) genes (temporal collinearity; [144]). The 3’ to 5’ activation is associated with corresponding change in epigenetic modifications from a repressive H3K27me3 modification to an activating H3K4me3 modification [145]. The dynamical change in histone modification is associated with a progressive shift in the 3D compartmental organisation of the cluster. Accordingly, when a *Hox* cluster is transcriptionally inactive (enriched in H3K27me3) it forms a single 3D compartment that can interact in *cis*- and *trans*-with distantly located loci that are also enriched in H3K27me3 [142,146]. As the 3’ to 5’ transcription of a *Hox* cluster proceeds there is a switch in 3D organization whereupon newly activated *Hox* genes beginning at the 3’ end of the cluster are progressively incorporated into a transcriptionally active compartment, while the rest remain in an inactive compartment [142]. By the time embryos exit the phylotypic progression temporal collinearity has established the spatially-restricted patterns of *Hox* gene expression (spatial collinearity; day 10.5 for the somitic *Hox* code; Figure 4C on the top), which are stable for the rest of development [147]. In a nucleus taken from the posterior trunk of the day 10.5 embryo much of the *Hox* cluster has taken up an “open”, euchromatic conformation that is permissive for gene expression (two blue “euchromatic” loops at the bottom right of Figure 4C) and only a small part of the *Hox* cluster remain assembled into a compact, silent Pc-G domain (one orange loop at the bottom right of Figure 4C). More anteriorly, around the mid-point of the trunk, only the 3’-most genes of the *Hox* gene cluster are in an open conformation (one blue “euchromatic” and two orange (Pc-G) loops at the middle on right of Figure 4C). In the forebrain where *Hox* genes are not expressed the entire *HoxD* cluster is assembled into a Pc-G domain (three orange loops at top on right in Figure 4C). Segregation of micro-phase separated Pc-G domains associated with the *Hox* genes contributes position-specific contact enrichments to ECDs (Diagram C in Box 1; cartoons on right of Figure 4C). Position-specific contacts mediated by Pc-G domains/complexes associated with *Hox* genes are but one fraction of the contacts that contribute to ECDs for, as explained below, there are around 2000 Pc-G domains/complexes in mouse and human genomes [148] all of which are likely to contribute to ECDs along with heterochromatin-*like* domains/complexes.

Temporal collinearity has been posited as the *cause* of the phylotypic progression - an ineluctable *Einbahnstraβe* [130] through which vertebrate embryos must pass in order for the spatially-restricted patterns to be established (spatial collinearity) thereby ensuring each position along the A-P has its own *Hox* code that is stably carried forward for the rest of development. We believe that the establishment of *Hox* gene spatial collinearity is taking place in the larger context of the deployment of heterochromatin-*like* domain/complexes during the phylotypic stage (Figure 4C on left). Together they contribute to the evolutionary restriction seen as the “bottleneck” in the hourglass model of vertebrate development (Figure 4A). Key questions remain. For example, what is the mechanism(s) by which the ground states of heterochromatin-*like* and Pc-G domains/complexes are achieved on days 8.25 and 6.5 respectively? A related question is: How is the *Einbahnstraβe* [130] converted into a two-way-street during the animal cloning procedure? The A-P axis must be faithfully recapitulated in reconstructed embryos even when the transferred nucleus retains the memory of the *Hox* code specific for only one position along that axis.

The number of Pc-G domains/complexes is around 2000 in the genomes of mouse and human ES cells, where there is a preference of PRC1 and PRC2 complexes to localise to CGIs [148]. The domains/complexes contribute to the coarse-grained chromatin-state pattern that characterises mammalian genomes [149–150]. That is, they contribute to the “segmented” nature of mammalian genomes, where segments or “blocks” consisting of Pc-G and heterochromatin-*like* domains/complexes alternate with segments or “blocks” of euchromatin. As shown (Diagram C in Box 1), ECDs are generated by *cis*- and *trans*-contacts between both types of micro-phase separated domains/complexes. How the theory of micro-phase separation of block copolymers (BCPs) might explain the behaviour of the domains/complexes as “blocks” in a chromatin fiber is the subject of the next section where, in the interests of clarity, we concern ourselves with heterochromatin-*like* domains/complexes although the same arguments apply to Pc-G domains/complexes.

## 5. Heterochromatin-*like* domains/complexes and block copolymers (BCPs)

Our survey of co-localisation of HP1s with H3K9me3 shows that depending on cell type there are (conservatively) around 163-855 heterochromatin-*like* domains (>0.1Mb) and 18853-32292 complexes (<0.1Mb) in the human genome (Table 1). The heterochromatin-*like* domains/complexes are contiguous with the nucleosome fiber and can be coarse-grained as alternating “blocks” of heterochromatin and euchromatin along the nucleosome fiber “polymer” (Box 1). This organisation is analogous to that of a BCP that contains a series of alternating blocks (*e.g*., A-type and B-type), each composed of multiple monomers (A monomers and B monomers). Where the monomers are incompatible the blocks segregate on the basis of like-with-like, with A-type blocks associating with A-type blocks and B-type associating with B-type. Accordingly, the BCP forms spatially segregated domains that are enriched in A or B [151]. We have drawn upon the basic polymer physics of micro-phase separation of bulk BCPs to explain how segregation of micro-phase separated heterochromatin-*like* domain/complexes is likely to generate contact enrichments that contribute to ECDs detected in Hi-C maps ([41]; Box 1). While we recognise that micro-phase separation seen with simple hydrocarbon BCPs (top of Box 2) is a useful analogy we argue in the next sections that the fundamental physics of micro-phase separation of heterochromatin-*like* domains/complexes is not the same as BCPs. In order to develop our argument we begin with a simplified description of the physics underpinning micro-phase separation of BCPs.

### Box 2: The Flory-Huggins parameter (χ) and the monomer as the “unit of incompatibility”.

Polymers are very long molecules formed from small repeating units called monomers. Polymers come in many varieties but the standard types are formed from small hydrophobic hydrocarbon monomers (~100Da) that have low ionisation and polarization potential:

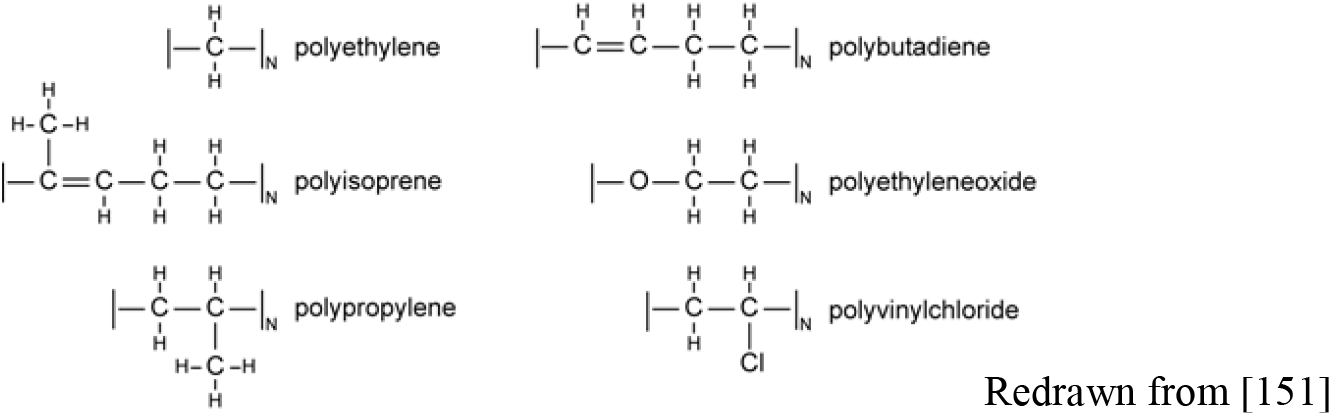

Homo-polymers formed from such monomers were the concern of Flory [154] and Huggins [155] when they set out to describe the thermodynamics of polymer mixing in simple solvents and, more importantly, the conditions under which they de-mix *i.e*., phase separate into polymer-rich and solvent-rich phases. For such homo-polymers the tendency to phase separate is described by the Flory-Huggins parameter χ (equation (1)), which quantifies the balance between the three types of interaction that take place when homo-polymers are added to a solvent, namely polymer-solvent, polymer-polymer and solvent-solvent interactions.

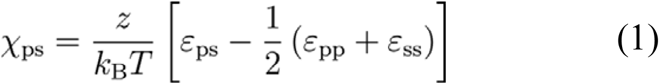

These interactions can be modelled using mean-field lattice theory where polymer and solvent molecules arrange themselves randomly within in an infinite lattice structure of coordination number z (see below for two-dimensional lattice where z = 8), each occupying one lattice position. The lattice is set at the volume occupied by one monomer segment of the polymer; it is assumed that the lattice is incompressible so the volume of the solvent is equivalent to that of one monomer unit:

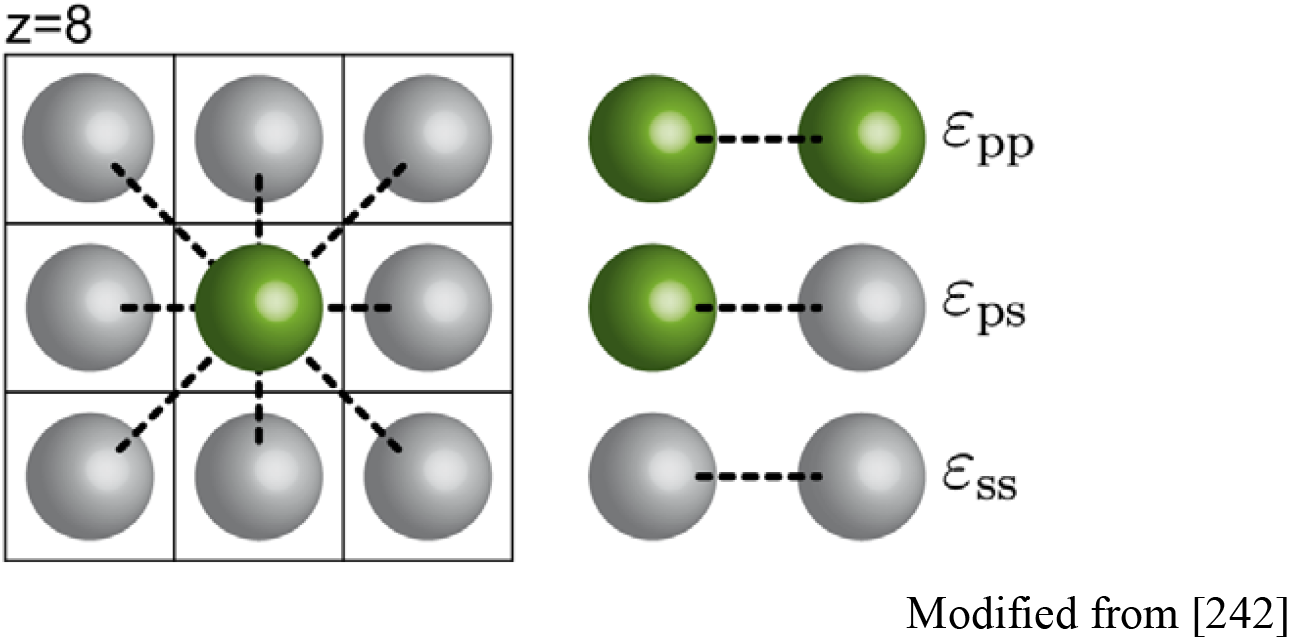

Whether the polymer phase separates or not depends on the mean field energies per lattice site, *i.e*., the energetic cost (in terms of thermal energy *k*_B_*T*) of having pairs of polymer-solvent monomer units (*ε*_ps_), pairs of polymer units (*ε*_pp_) and pairs of solvent units (*ε*_ss_) next to each other in the lattice. In the case of highly unfavourable interactions, where there is an energetic cost for having the polymer monomer site adjacent to the solvent site, monomers will prefer to be near each other (like-with-like) and phase separation is likely to occur (poor solvent; χ>0). Above a particular value, χ_crit_, the critical value of the Flory-Huggins parameter where the energy of interaction overcomes the entropy of mixing, the polymer will collapse and phase separation is observed (see [242] for a more detailed discussion of the Flory-Huggins free energy of mixing equation). χ< 0 in the case of highly favourable interactions between monomer and solvent, where monomers will avoid being near each other whereupon the polymer chain swells (good solvent). Thus χ is a measure of the degree of incompatibility of the polymer with the solvent and, as defined in equation (1), is inversely proportional to temperature. The sign of χ (+ve or −ve) is dependent upon the choice of monomer repeating unit; the “*unit of incompatibility*” for the Flory-Huggins approach to the miscibility of polymers is the *monomer*.

BCPs can be configured into a wide variety of molecular architectures based on two, three or more monomer types [152]. Of these architectures, symmetric BCPs containing equal sized blocks A and B (AB di-BCPs) have been the focus of a very large number theoretical and experimental studies and our discussion of BCPs will deal almost exclusively with this simplest form of BCP. The physics of the phase behaviour of a bulk (undiluted) AB di-BCP centres upon the covalent bond that separates the two chemically dissimilar blocks and prevents *macroscopic* phase separation (for a review see [153]). The bond makes the entropy of mixing small and excess free energy generated by even minor chemical or structural differences between A and B blocks is sufficient to produce contributions that are unfavourable to mixing. Put another way, phase separation of a bulk di-BCP is driven by an unfavourable mixing enthalpy coupled with small mixing entropy, with the covalent bond connecting the blocks preventing macroscopic phase separation. *Microscopic* phase separation of a di-BCP depends on three parameters [153]: (1) the volume fractions of the A and B blocks (*f_A_* + *f_B_* = 1), (2) the total degree of polymerization (*N* = *N_A_* + *N_B_*), and (3) the Flory–Huggins parameter (χ_AB_). The χ-parameter specifies the degree of incompatibility between the A and B blocks and this is what ultimately drives micro-phase separation. The relationship between χ_AB_ and temperature (T) is given in equation (2), which is an application to BCPs of the original mean field lattice theory [154,155] for the thermodynamic behaviour of homo-polymers in a simple solvent (see equation (1) in Box 2):

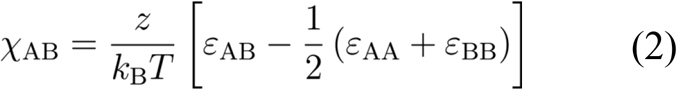

Equation (2) describes the energy cost (in units of thermal energy *k_B_T*) when A and B monomers make contact. z is the coordination number of an incompressible lattice (Box 2) and represents the number of nearest neighbours per lattice site that can be occupied by either an A-type monomer or a B-type monomer. ε_AB_, ε_AA_, and ε_BB_ are the contact energies per repeat unit (monomer) of A–B, A–A, and B–B, respectively. As with equation (1) in Box 2 the sign (positive or negative) and magnitude of χ_AB_ is determined by the choice of monomer – the “*unit of incompatibility*” is the monomer. For a typical di-BCP consisting of two types of simple hydrocarbon monomer, such as polyisoprene-block-polystyrene (PI-b-PS), where electrostatic interactions are negligible (*i.e*., governed by dispersive interactions), the value for χ is positive and small (~0.1). The *positive* value indicates that there is a net repulsion between the PI and PS blocks of the PI-b-PS and they have a tendency to micro-phase separate; a di-BCP that has a *negative* value of χ would indicate a free-energy drive towards mixing. As seen in equations (1) and (2) χ varies inversely with temperature. Increasing temperature or decreasing χ_AB_ through choice of monomers reduces the incompatibility between the constituent blocks and combinatorial entropy increases, resulting in mixing whereupon the di-BCP becomes disordered (*i.e*. homogeneous).

We have previously drawn on the observation that χ varies inversely with temperature and that micro-phase separation is dependent upon composition (volume fraction), to provide insight into how changes in the activity of chromatin-associated cohesin affects the compartmentalisation observed in Hi-C maps (Figure 5) [41]. As explained ([41]; Diagram B in Box 1; see also sections 7 and 8 below), the incompatibility between the heterochromatin-*like* domains/complexes and euchromatin is owing to the “bridging” of the H3K9me3-marked nucleosomes and it is this bridging that drives micro-phase separation of the domains/complexes from euchromatin. Far *cis*- and *trans*-contacts between micro-phase separated heterochromatin-*like* complexes results in segregation and contributes to the emergence of the ECDs observed in Hi-C maps ([41]; Diagrams B and C Box 1). Mixing of heterochromatin-*like* complexes with euchromatin is caused by cohesin, which is a loop-extruding factor (LEF) [156,157]. LEFs attach to the chromatin fiber and reel it in from both sides, thereby extruding a progressively growing chromatin loop until the LEFs fall off, bump into each other, or bump into extrusion barriers such as CTCF, which define TAD boundaries [156–158]. Loop extrusion is an energy-driven, ATP-dependent, process [159]. Mixing is promoted by friction of the heterochromatin-*like* domain/complex with the nucleoplasm during loop extrusion, which converts the kinetic energy of loop extrusion into thermal energy (work done by ATP-hydrolysis is converted into heat). As a consequence, HP1-mediated “bridging” of H3K9me3-marked nucleosomes is disrupted making the domains/complexes less “heterochromatic” and more “euchromatic”, with the smaller domains/complexes undergoing greater mixing compared to the larger domains/complexes in keeping with the dependency of micro-phase separation on volume fraction.

**Figure 5:**
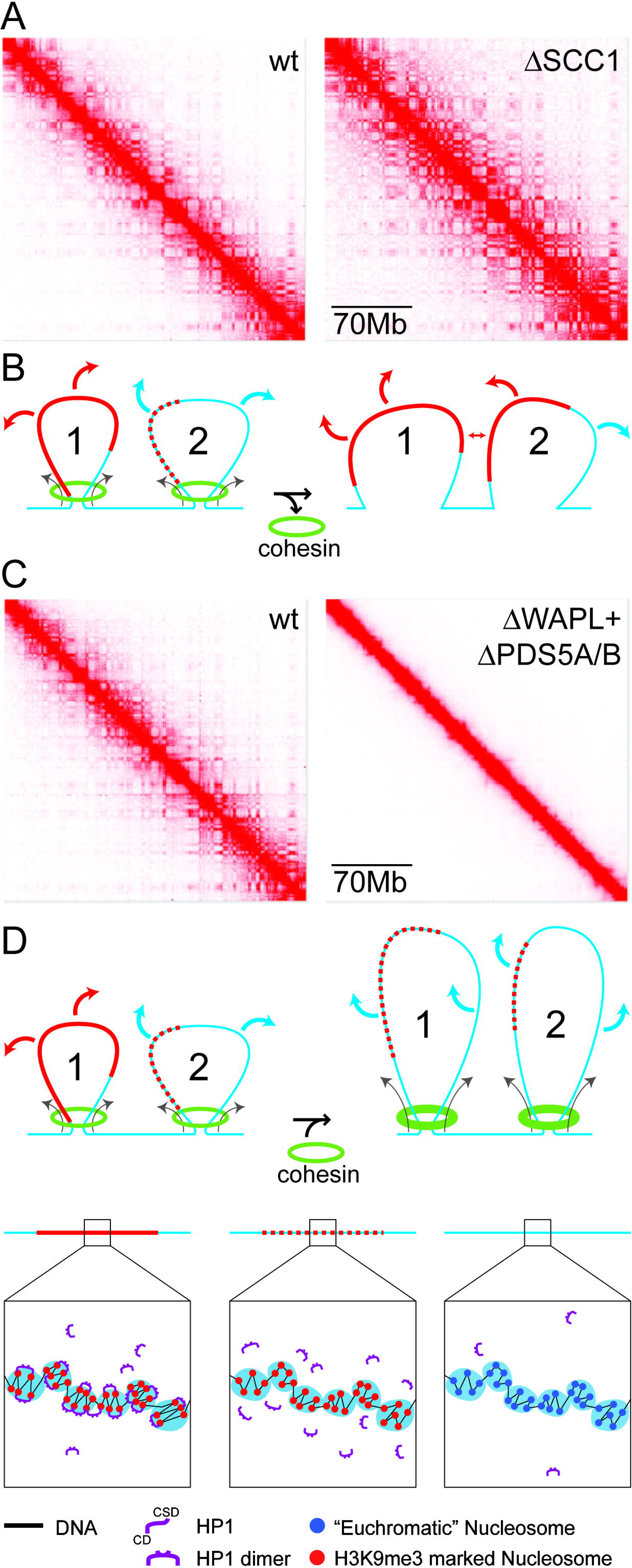
Mixing of heterochromatin-like domains/complexes with euchromatin by modulating chromatin-associated cohesin activity. **(A) Hi-C maps for wt and SCC1 depleted cells at low reso1ution (70Mb) to show the effect of cohesin depletion on compartmentalisation.** The Hi-C map generated from SCC1 deleted cells (on right) have a finer, better defined, compartmentalisation compared to the Hi-C map from wt cells (on left). Taken and modified from [161], with permission. **(B) A BCP-based model that provides an explanation for the effect of SCC1 depletion on compartmentalisation**. On the left are two loops that are being extruded by chromatin-associated cohesin LEF. Loop 1 contains a large micro-phase separated heterochromatin domain/complex (continuous red line) that is flanked by euchromatin (continuous blue line). The large domain/complex is resistant to mixing; bridging of H3K9me3-marked nucleosomes by HP1 is not disrupted (see key at bottom of figure). Loop 2 contains a smaller domain/complex (smaller “block”) that is subject to mixing (dotted line; see key at bottom of figure) by loop extrusion, which is a prediction of BCP theory, where a block of smaller volume fraction has greater tendency for mixing, with the BCP becoming “homogeneous” (one phase). Loop 1 makes far *cis*- (not shown) and *trans*-contacts (red arrows) with other HP1-containing domains/complexes which result in contact enrichments that emerge in ECDs. Loop 2 makes euchromatic contacts (blue arrows) that will fall into A-type compartments because the smaller domain/complex is now more euchromatic and less heterochromatic as a consequence of mixing. On the right are the same two loops after chromatin-associated cohesin is eliminated by loss of SCC1, whereupon loop extrusion ceases and mixing is reduced. This has little effect on the larger domain/complex in loop 1, but leads to the reconstitution of the smaller domain/complex in loop 2 (now a continuous red line) as a consequence of bridging H3K9me3-marked nucleosomes by HP1. The newly-reconstituted domain/complex can then make far *cis*- (double-headed arrow) and *trans*-contacts (red arrows) that result in the finer, more defined, compartmentalisation seen Hi-C maps from SCC1 deleted cells. **(C) Hi-C maps for wt and WAPL/PDS5A/B depleted cells at low reso1ution (70Mb) to show effect of enhancing cohesin activity on compartmentalisation.** The Hi-C map generated from WAPL/PDS5A/B deleted cells (on right) where cohesin activity is enhanced there is little or compartmentalisation compared to the Hi-C map from wt cells (on left). Taken and modified from [161], with permission. **(D) A BCP-based model that provides an explanation for the effect of WAPL/PDS5A/B depletion on compartmentalisation.** The same two loops, loops 1 and 2, as seen on the left in **(B)**, where loop extrusion by chromatin-associated cohesin leaves the larger heterochromatin-*like* domain/complex in loop1intact, but the smaller domain/complex in loop 2 undergoes mixing. As before, in wt cells, loop 1 makes far *cis*- (not shown) and *trans*-contacts (red arrows) with other domains/complexes which result in contact enrichments that emerge in ECDs. Loop 2 makes euchromatic contacts (blue arrows) that will fall into A-type compartments. In WAPL/PDS5A/B depleted cells the activity of cohesin is *enhanced* (thicker green circles at base of loops). The effect on compartmentalisation is striking. Compartmentalisation is almost completely eliminated. One cause of this loss is that increased loop extrusion results in dissolution of the larger (loop 1) and smaller (loop 2) domains/complexes (dotted lines; see key at bottom of the figure) and mixing with euchromatin. If this occurs genome-wide there would be a loss of compartmentalisation. As shown, loop 1 and 2 make *cis*- (not shown) and *trans*-contacts (blue arrows) with other euchromatic chromatin fibers, however, continued extrusion eventually leads to collapse of interphase chromatin organisation, whereupon chromatin takes on condensed mitotic-like chromatin state termed vermicelli [169]; it seems likely that compartmentalisation would also be affected by this loss of organisation. The “clutch” diagrams were modified from [41].

The above provides a framework for understanding how mutants that affect cohesin activity can in turn affect ECDs observed in Hi-C maps (Figure 5). For example, when the cohesin subunit SCC1 is deleted [160] finer, better defined, ECDs emerge in Hi-C maps (Figure 5A, on the right; [161]) that are normally “masked” in wild-type (wt) cells (Figure 5A, on the left). The emergence of the finer compartmentalisation in SCC1 mutants can be understood in terms of the model in Figure 5B, that consists of two loops, loops 1 and 2. Loop 1 contains a large micro-phase separated heterochromatin-*like* domain. In loop 2 is a smaller heterochromatin-*like* complex. In wt cells, extrusion of the large heterochromatin-*like* domain in loop 1 by the cohesin complex (green ring on left in Figure 5B) has little effect on the mixing of the domain with euchromatin; the domain makes far *cis*- and *trans*-contacts (red arrows) that will be detected as contact enrichments in ECDs. Extrusion of the smaller complex in loop 2 by the cohesin complex results in extensive mixing with euchromatin leading to its dissolution in wt cells (depicted by the red dots on blue line). As a consequence, the smaller complex makes contacts (blue arrows) that will be detected as contact enrichments in A-type compartments. On the right in Figure 5B is an explanation of the emergence of the finer compartmentalisation in SCC1 deleted cells. Here energy-driven loop extrusion is absent and the HP1-mediated bridging of H3K9me3-marked nucleosomes in the small heterochromatin-*like* complex is reconstituted (red line in loop 2 on right of Figure 5B); the large domain that precedes it is unaffected. Reconstitution of the small complex promotes far *cis*- and *trans*-contacts (red arrows) that cause the finer, more defined, ECDs observed in SCC1 deleted cells (Figure 5A on right).

Mixing of larger heterochromatin-*like* domains with euchromatin by *enhanced* energy-driven loop extrusion could go some way to explaining the changes in compartmentalisation seen in Hi-C maps derived from WAPL/Pds5A/B compound mutant cells (Figure 5C on the right; [161]). By way of background, WAPL normally removes cohesion through binding of its YSR motifs to the regulatory subunit of cohesin, Pds5 [162,163]. The WAPL-pds5 interaction stabilizes a transient, open, state of the cohesin ring that results from disruption of the interface between the SMC3 and Rad21/Scc1 cohesin subunits, with consequent release of the sister chromatids [164,165]. In mutations or “knockdowns” of WAPL cohesin is retained along the length of the chromosome arms [166–168]. Retention of cohesin complexes results in continued loop extrusion, the formation of larger topologically associated domains (TADs) and the loss of compartmentalisation [169] (Figure 5C, on the right). As shown in Figure 5D (on the right), loss of compartmentalisation may, at least in part, be explained by enhanced loop extrusion that disrupts HP1-mediated bridging of H3K9me3-marked nucleosomes even within large heterochromatin-*like* domains (red dots on blue line) resulting in mixing of domains/complexes with euchromatin. Unrestrained loop extrusion eventually leads to an overall loss of interphase chromatin organisation where interphase chromatin takes on condensed mitotic-like chromatin state termed vermicelli [169]. This loss of interphase organisation will also affect compartmentalisation.

Recent work in fission yeast may provide additional support for loop extrusion as a mechanism for mixing of heterochromatin-*like* domain/complexes with euchromatin. Specifically, it was observed that the *Pds5* mutation in fission yeast alleviates heterochromatin-mediated silencing at the donor mating-type region [170]. Notably, the effect of the *Pds5* mutation could be reversed by introduction of the *eso1* mutation [170]. eso1p is an acetyltransferase required for establishing stable cohesin complexes on chromatin, which it does by acetylating the heads of the SMC3 cohesin subunit that protects against removal of cohesin by WAPL [171,172]. Genetic interaction of the two mutations can be explained by reversible mixing of the heterochromatin-*like* complex assembled at the silenced mating-type region with flanking euchromatin. Accordingly, loss of Pds5p activity enhances cohesin-driven loop extrusion leading to disruption of HP1-mediated bridging of H3K9me3-marked nucleosomes resulting in mixing with euchromatin (*c.f*. Figures 5C and D on right). Introduction of *eso1* returns cohesin activity to near normal levels by interrupting cohesin establishment and that, in turn, reverses mixing and reconstitutes the heterochromatin-*like* complex and silencing at the donor mating-type region.

Micro-phase separation of a BCP is dependent upon the sign and magnitude of χ, *i.e*., the degree of incompatibility of the monomer (“unit of incompatibility”; equation (2); Box 2) repeating units that make up the blocks. That BCP theory could provide an explanation for how cohesin activity might regulate mixing of heterochromatin-*like* domains/complexes with euchromatin (Figure 5; [41]) indicates that the same is true for heterochromatin-*like* and euchromatic “blocks” in the chromatin fiber. Put another way, there exists a “unit of incompatibility” that is the repeating unit from which heterochromatin-*like* and euchromatic “blocks” are assembled and it is the degree of incompatibility between these heterochromatin-*like* and euchromatic repeating units, specified by χ, which drives micro-phase separation of heterochromatin-*like* domains/complexes from euchromatin. Notably, virtually all modern theories of micro-phase separation employ this simple one-parameter thermodynamic description of the driving force for micro-phase separation [173,174]. Fortified by these credentials, we move to exploring whether the approach taken for the determination of χ for di-BCPs can also be used to estimate χ for heterochromatin-*like* domains/complexes flanked by euchromatin.

## 6. Di-BCPs, χ and the monomer as the “unit of incompatibility”

For di-BCPs, χ can be described by (2) where interactions between the hydrocarbon *monomer* repeating units (see top of box 2) are dominated by dispersive (van der Waals) interactions. In a classical thermodynamic approach for determination of χ based on these dispersive interactions, Bates [175] showed that the contact energy (*ε_ij_* between *i* and *j segments*, each consisting of (dissimilar) monomer repeating units, could be quantified by the equation:

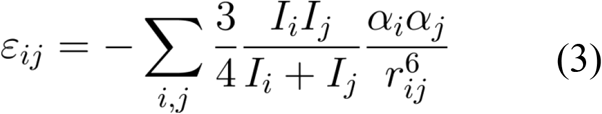

*r_ij_* is the segment-segment separation, *α* and *I* are the segment polarizability and the first ionization potential, respectively. If there is neither a change in volume (equation (2) assumes an incompressible lattice; see also Box 2) nor a preference for a particular segment orientation upon mixing, passing the binary interaction energies (for A-A, B-B and A-B) quantified in (3) through equation (2) gives the following result for χ:

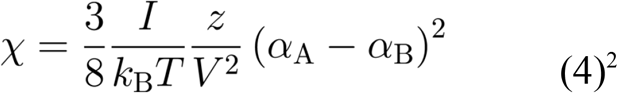

Equation (4) is an approximation for χ owing to a number of assumptions (see [175]; FS Bates, personal communication). A cubic lattice is assumed where *I_j_* = *I_i_* ≡ *I*; the latter is valid to within 10% for most hydrocarbons in a di-BCP. The volume of the cubic lattice site is also assumed as *V*, where *V* = *r^3^* (the *r^6^* term is represented as *V^2^* in (4)). Further, defining the number of segments that interact with each other by the co-ordination number z places all segments surrounding a particular segment within a single parameter, with an average interaction given by *I* and (α_A_ - α_B_)^2^. A more rigorous treatment that removes the assumption of an incompressible (cubic) lattice [154,155] would involve a calculation based upon summation over all neighbouring segments with the r^6^ dependence (FS Bates, personal communication). Despite being an approximation equation (4) provides qualitative predictions, for example, for a bulk di-BCP governed solely by dispersive interactions a value of χ ≥ 0 is obtained indicating that the di-BCP will have a tendency to phase separate [175]. In this (classical) treatment the magnitude of χ is fundamentally dependent upon the choice of repeating unit, the monomer. The monomer is the *unit of incompatibility* from which the “blocks” or “segments” of a di-BCP are made.

As might be expected, the assumptions required to derive (4) are rarely satisfied in practice. Because of this, χ is usually determined empirically using a much simpler general formula:

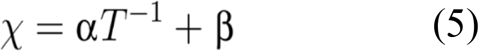

χ retains the temperature dependency. α and β are experimentally determined coefficients for enthalpy and excess entropy for a particular composition of a BCP. We believe that the utility of equation (5) can be extended to enable an estimation of χ for a heterochromatin-*like* domain/complex vs euchromatin once the *unit of incompatibility* has been defined. And this is the subject of our next section.

## 7. The oligo-nucleosomal “clutch” as the “unit of incompatibility” of chromatin

Estimation of χ for heterochromatin-*like* domains/complexes flanked by euchromatin is likely to employ a different physics to that which developed for determination of χ for di-BCPs (equations (2) to (4)). An obvious reason for this is that heterochromatin-*like* domains/complexes are assembled from the 11nm “beads-on-a-string” nucleosome fiber “polymer” that is the primary structure of chromatin [1]. The “bead”, or *monomer* repeating unit, is the nucleosome, a multicomponent structure very different to the simple hydrocarbon *monomer* found in BCPs (see top of Box 2). By comparison the nucleosome is gigantic. The molecular weight of the nucleosome core particle, consisting of 147bp of DNA that is wrapped 1.75 times in left-handed super-helical turns around a histone octamer, is over three orders of magnitude greater (~200kDa; [176]) than the hydrocarbon monomers of BCPs (~100Da; top of Box 2). In addition, monomers in BCPs are hydrophobic and dispersive interactions dominate (equation (3)), while the nucleosome is a highly electrostatic multicomponent structure. A single histone octamer in the nucleosome has ~220 positively charged lysine and arginine residues and ~74 negatively charged aspartic acid and glutamic acid residues. The phosphate backbone of 200bp of DNA that includes the 147bp associated with the nucleosome core particle plus linker DNA adds a further 400 negative charges [177]. Given the obvious differences in physiochemical properties it is a wonder that BCP theory could be used as an analogy to understand how ECDs might emerge by micro-phase separation of heterochromatin-*like* complexes (Figure 5; [41]). This most likely has deep roots in their shared polymeric nature [178,179] but does not mean that the physics of micro-phase separation and segregation of heterochromatin-*like* domains/complexes and BCPs is the same. Nor may we expect that the degree of incompatibility, as designated by χ, is caused by the same physiochemical mechanisms.

For di-BCPs the sign and magnitude of χ is determined by the degree of incompatibility of the two dissimilar monomers; the monomer is the “*unit of incompatibility*” from which the microphase separated “blocks” or “segments” are composed (Box 2; equations (2) and (4)). For heterochromatin-*like* domains/complexes the sign and magnitude of χ will be determined by the degree of incompatibility of the domains/complexes with euchromatin. We suggest that incompatibility between domains/complexes and euchromatin is the result of excess free energy generated by binding of HP1 to H3K9me3-marked nucleosomes in the 11nm nucleosome fiber combined with excess entropy that results from compaction of the nucleosome fiber by the “bridging” effect [29,180,181]. This leads to an unfavourable enthalpy of mixing of the heterochromatin-*like* domain/complex with euchromatin. This begs the question: what is the “unit of incompatibility” for a heterochromatin-*like* domain/complex? On the face of it, it would seem that the unit of incompatibility is the H3K9me3-marked nucleosome *monomer* in the same way that the hydrocarbon *monomer* is the unit of incompatibility for di-BCPs. However, experimental and theoretical work indicates that: (i) HP1 proteins drive the incompatibility rather than the H3K9me3-marked nucleosomes *per se* and, (ii), the unit of incompatibility is a larger than the mononucleosome. For one, recent liquid Hi-C experiments directed towards identifying factors that generate and maintain compartmental domains indicated that ECDs (B-type compartmental domains) enriched in HP1α and β were found to be equally stable after chromatin fragmentation followed by HP1γ, with ECDs enriched in the Polycomb CBX8 homologue being the most unstable [182]. K9me3 (to which HP1α and HP1β bind) is essentially indistinguishable compared to K27me3 (to which Polycomb CBX homologues bind) *i.e*., K9me3 and K27me3 are unlikely to contribute to differences in chemical potential between nucleosomes possessing these modifications, especially given the highly electrostatic environment of the nucleosome (see discussion above). Yet there are great differences in the dissociation kinetics of ECDs enriched in HP1 a and β compared to CBX8 indicating that it is the “bridging” protein that drives micro-phase separation and generation of ECDs rather than the histone modifications *per se*. Second, theoretical work using simple models where binding proteins (“binders”) cross-link polymer-specific binding sites have been able to recapitulate many features of Hi-C maps where binders cause folding of the polymer and phase transitions [183,184]. The binding sites on the polymer are neutral with respect to phase transitions and serve simply as binding sites for the “binders” that are the actual drivers of the transitions. Polymer simulations using a similar model have shown that it is HP1 binding to H3K9me3-marked nucleosomes that drives phase separation with the minimum H3K9me3-marked segment that can be phase separated by HP1 “bridging” being around 20kb [185]. Finally, compaction of heterochromatin-like domains/complexes requires at least two H3K9me3-marked nucleosomes to be “bridged” by HP1 ([181]; Box 1). This indicates that the “unit of incompatibility” is more than a single H3K9me3-marked nucleosome: it is not a monomer. It is likely to be larger than two nucleosomes because *in vitro* studies show HP1-mediated bridging can compact nucleosome arrays into clusters of nucleosomes [29,180]. Similar observations have been made with Polycomb CBX proteins where, for example, the CBX2 protein, as part of the PRC1 complex, has intrinsic chromatin compaction activity where it is able to compact at least four nucleosomes ([186,187]; Diagram A in Box 1, bottom row). The unit of incompatibility is likely to consist of a cluster or “clutch” of nucleosomes.

There is a weight of experimental evidence showing that “clutches” of 2 to 10 nucleosomes with variable degrees of zig-zag geometry are a ubiquitous motif within interphase chromatin; this organisation may represent the secondary structure of interphase chromatin [188]. Clear evidence came from super reso1ution microscopy, which showed that chromatin outside constitutive heterochromatin was characterized by the assembly of irregularly folded ‘clutches’ containing 2 to10 nucleosomes while the density of larger ‘clutches’ was greater within constitutive heterochromatin [189]. A variety of electron microscopy approaches have confirmed the presence of small clumps of 2 to 10 nucleosomes *in vivo* without any evidence for longer stretches of organized nucleosome fiber [190–193]. Nucleosomes in the “clutches” possess, to a lesser or greater extent, a local zig-zag organisation. Radiation-induced spatially correlated cleavage of DNA with sequencing revealed zig-zag geometry of short stretches of the nucleosome that was noticeably enriched in H3K9me3-marked nucleosome fibers in constitutive heterochromatin [194]. *In vivo* studies using controlled DNA breakages [195] or cross-linking of nucleosomes to one another followed by digestion and electron microscopy [196] have also given results consistent with there being short stretches of nucleosomes (3-10 nucleosomes) with zig-zag geometry in the nucleus. A substantial number of i*n vitro* studies using synthetic nucleosomal templates have been confirmatory in nature and revealed a zig-zag motif for short stretches of (4 to 12) nucleosomes [197–199]. Notably, it is has been shown in a wide variety of eukaryotes that nucleosomes are connected by linkers biased towards non-integer DNA double-helical turns (*e.g*., 0.5, 1.5, 2.5 turns) [200]. Such fibers possess zig-zag nucleosomal geometry [201] and exhibit enhanced phase separation [202].

Based on the evidence we suggest that the “*unit of incompatibility*” for chromatin is a dynamic multi-component structure unlike the indivisible hydrocarbon monomer (top of Box 2) that is the unit of incompatibility for BCPs. Specifically, for euchromatin the unit of incompatibility consists of an oligo-nucleosomal “clutch” of 2 to 10 nucleosomes where the nucleosomes within the “clutch” are disorganised with weak zig zag geometry (Figures 6B and C). For heterochromatin-*like* domains/complexes the unit of incompatibility is an oligo-nucleosomal “clutch” of 2 to 10 H3K9me3-marked nucleosomes “bridged” by HP1 dimers that compact and stabilize the zig-zag geometry of nucleosomes within the “clutch” (Figures 6B and C). It is the incompatibility between the dissimilar repeating units (euchromatic “clutch” vs heterochromatin-*like* “clutch”) that determines the magnitude of χ. This value will indicate whether a there is a tendency for a heterochromatin-like domain/complex to micro-phase separate from the surrounding euchromatin. We next describe a theoretical “clutch” model for the determination of χ for a heterochromatin-*like* “clutch” vs euchromatic “clutch” based on equation (5).

**Figure 6:**
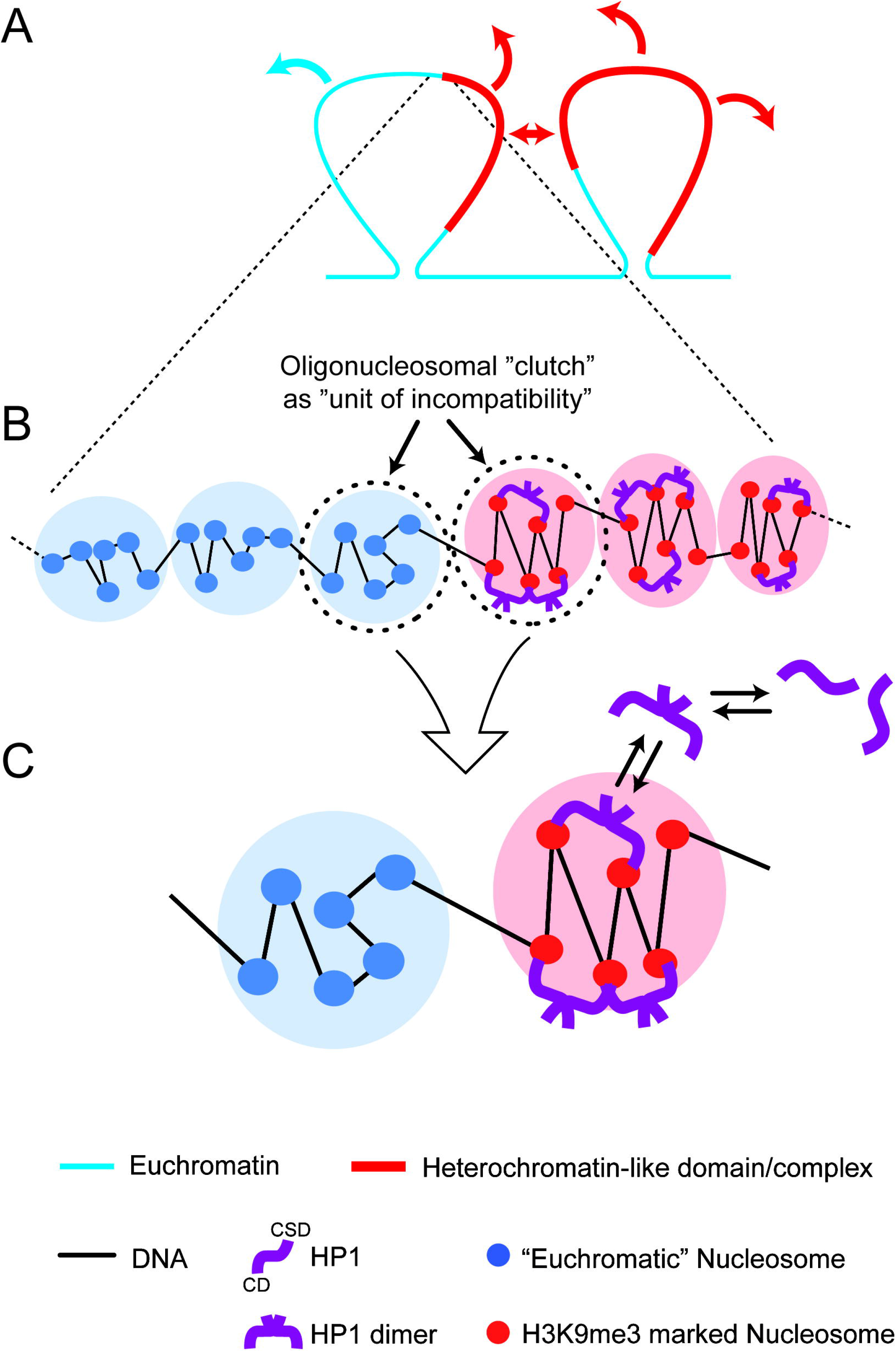
The oligo-nucleosomal “clutch” as the “unit of incompatibility” of chromatin. **(A) Chromatin loops containing two heterochromatin-*like* domains/complexes (red line) flanked by euchromatin (blue line).** Heterochromatin-*like* domains/complexes make *cis*- (doubled-headed arrow) and *trans*- (red arrows) contacts other micro-phase separated heterochromatin-*like* domains/complexes (Diagram B in Box 1). The *cis*- and *trans*-contacts between micro-phase separated heterochromatin-*like* domains/complexes generate contact enrichments seen in ECDs (Diagram C in Box 1). The euchromatic segments can also make *cis*- (not shown) and *trans*- (blue arrow) contacts with other euchromatic segments that are detected as A-type compartments in Hi-C maps. **(B) A region of the loop in (A) is magnified (dotted lines) to detail the junction between a heterochromatin-*like* domain/complex and a euchromatic segment.** The heterochromatin-*like* domain/complex is made up of “clutches” (pink circles), where each clutch contains 6 H3K9me3-marked nucleosomes. The H3K9me3-marked nucleosomes are “bridged” by HP1 dimers where the nucleosomes have distinct zig-zag geometry; a clutch so organised is the “unit of incompatibility” of the heterochromatin-*like* complex. The “unit of incompatibility” of the euchromatic segment are “clutches” also composed of 6 nucleosomes where the nucleosomes are more disorganised with only weak zig-zag geometry. **(C) Incompatibility of the euchromatic “clutch” vs heterochromatin “clutch” is caused by HP1 bridging of H3K9me3-marked nucleosomes.** This depicts the theoretical model (see section 8 and equation (6)). Absent HP1 the heterochromatin-*like* clutch containing H3K9me3-marked nucleosomes is thermodynamically equivalent to the euchromatic clutch. Incompatibility is caused by HP1 bridging of H3K9me3-marked nucleosomes. Monomers of HP1 dimerise and the dimers “bridge” H3K9me3 marked nucleosomes, which has the effect of stabilising the zig-zag geometry and compacting the nucleosomes within the clutch. This situation can be compared to that of di-BCPs where incompatibility between monomers is driven by excess free energy owing to small chemical and/or structural differences, whereas for heterochromatin-*like* domains/complexes incompatibility between “clutches” is driven by excess free energy that results from bridging of H3K9me3-marked nucleosomes by HP1. “Clutch” diagrams were modified from [41].

## 8. Theoretical “clutch” model

As shown (Figure 6C) the model consists of the following:

1. A symmetric pair of “clutches”, a euchromatic “clutch” and heterochromatin-like “clutch” that are connected by linker DNA. Each clutch contains 6 nucleosomes, a value that lies in the middle of the range 2 to 10. The heterochromatic-*like* “clutch” consisting of HeK9me3-marked nucleosomes represents the repeating unit (“*unit of incompatibility*”) of a heterochromatin-*like* domain/complex; likewise for the euchromatic clutch and euchromatin. Both clutches are equally miscible in the nucleoplasm (a concentrated solution of chromatin fibers) reducing the problem to incompatibility between the clutches.
2. In the absence of HP1, the “clutches” are thermodynamically equivalent. Incompatibility between the heterochromatin-*like* “clutch” (HC) and euchromatic “clutch” results from “bridging” of two K9me3 motifs on histone H3 proteins in separate nucleosomes by HP1 dimers; stacked nucleosomes are the preferred HP1 binding site. Bridging is maintained by constant exchange of bound HP1 dimers with free dimers in the nucleoplasm.
3. HP1 “bridging” stabilizes the zig-zag geometry of H3K9me3-marked nucleosomes and leads to compaction. Nucleosomes in the euchromatic “clutch” are disorganised with limited (unstable) zig-zag geometry.

Using equation (5), χ for the HC vs euchromatic “clutch” can be written as:

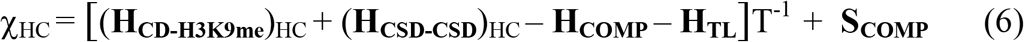

**H_CD-H3K9me_** is the free energy contribution from the binding of the HP1 CD to H3K9me3-marked histone H3. CD-H3K9me3 binding is best understood for HP1β [29,35] (Figure 1). Initial binding is *via* a non-specific electrostatic interaction of the N-terminal extension of HP1β with the H3K9me3 tail [29]. This causes the CD N-terminal region to draw upwards and wrap around the peptide and, as a consequence, an aromatic “cage” is formed from three conserved aromatic residues: Tyr21, Trp42 and Phe45 (Figure 1B). The majority free energy contribution comes from the electrostatic cation–π interactions where the positively charged (cation) methyl ammonium moiety is attracted to the negative electrostatic potential of the aromatic groups’ π-system [36]. The HC sub-script indicates that it is the sum of CD-H3K9me3 interactions that “bridge” different nucleosomes within the clutch that gives the total free energy contribution.

**H_CSD-CSD_** is the free energy contribution from the dimerization of the HP1 CSD. It is the dimeric form of HP1 that “bridges” H3K9me3-marked nucleosomes and drive micro-phase separation. The HP1β CSD forms a tight homodimer and the (CSD-CSD) interaction is of high affinity with sedimentation analysis indicating that the upper limit for the dissociation constant as <150 nM [203]. The HC sub-script indicates that it is the sum of CSD-CSD dimers that mediate “bridging” interactions between different nucleosomes within the clutch that gives the total free energy contribution.

**H_COMP_** is potential energy owing to: (i) elasticity of linker DNA (*i.e*., resistance to bending, twisting, and stretching deformation) and, (ii), steric exclusion between the nucleosomes and linker DNA [204]. H_COMP_ is a consequence of the “bridging” of H3K9me3-marked nucleosomes by HP1 dimers, which stabilises the zig-zag geometry and promotes compaction of nucleosomes in the “clutch”.

**H_TL_** is elastic potential energy of the terminal linker DNA (*i.e*., resistance to bending, twisting, and stretching deformation) that connects a heterochromatic-*like* clutch to a euchromatic clutch. For a heterochromatin-*like* domain/complex flanked by euchromatic segments the term would be 2**H_TL_**.

**S_COMP_** is the excess entropy given up to the nucleoplasm after “bridging” of H3K9me3-marked nucleosomes by HP1 dimers, which stabilises the zig-zag geometry and promotes compaction.

### 8.1 Characteristics and caveats of the theoretical “clutch” model

1. In the form presented the model may not be applicable to *macro*-phase separation of cytologically-visible constitutive heterochromatin. There are additional factors that may need to be included that are together severally necessary and jointly sufficient in causing macro-phase separation of constitutive heterochromatin. They include repetitive DNAs, ncRNAs, and proteins that bind (modified) DNA/histones [205]. There is also a documented interaction of Swi6^HP1^ CSD dimer with the H2Bα1 helix that results in the deformation of the nucleosome core that is thought to drive phase separation of constitutive heterochromatin in fission yeast [53]. These factors are not included in our model since we are concerned with micro-phase separation of heterochromatin-*like* domains/complexes *outside* constitutive heterochromatin.
2. H3K9me3 epigenetic modification acts as an HP1 binding site only. Absent HP1 a H3K9me3-marked “clutch” is thermodynamically equivalent to the euchromatic “clutch”. Incompatibility that increases the tendency to micro-phase separate is a consequence of HP1 “bridging” between H3K9me3-marked nucleosomes. Experimentally, a reference “clutch” consisting of 6 H3K9me3-marked nucleosomes could be used to measure changes in free energy and excess entropy generated after addition of HP1 to the system. The H3K9me3-marked clutch would need to be tethered to mimic the connection through linker DNA to flanking euchromatic “clutches”.
3. HP1dimer binding is modelled as “bridging” two H3K9-methylated histone H3 molecules in separate nucleosomes [37]. Accordingly, a nucleosome chain with stacked nucleosomes would be the preferred HP1 binding sites [206], which would be consistent with HP1 binding to a fiber with nucleosomes having a zig-zag geometry. Compaction would be promoted by allosteric cooperativity arising from changes in di-nucleosome conformation after HP1 bridging that would in turn enhance further HP1 bridging [206]. No direct HP1-HP1 co-operative binding is assumed, although there evidence that such cooperative binding might contribute to “spreading” of the heterochromatic state [207,208]. This has been modelled previously [204] and could be incorporated as a free energy contribution (**H**_HP1-HP1_)_HC_ to equation (6).
4. Histone acetylation is not included in the model. Histone acetylation is an epigenetic modification generally associated with euchromatin and is known to neutralize the positive charge on the histone tails [209]. Inclusion of histone acetylation could affect the incompatibility between the euchromatic and heterochromatin-like clutches. Our reasoning for not including histone acetylation is that H3K9me3 and H3K27me3 are truly epigenetic and define ECDs (Box 1). They are associated with “*write-and-read*” activities that ensure their inheritance from one cell generation to the next [210]. Histone acetylation does not possess an associated write-and-read activity.
5. The **H_TL_** term reduces the magnitude of χ and acts against phase separation. Accordingly, release of a domain/complex from the constraints of the chromatin fiber by severing linker DNA that connects the domain/complex to flanking euchromatin would remove the HTL term and enhance phase separation. Liquid Hi-C experiments have provided evidence that this is the case: chromatin fragmentation that releases domains/complexes from the chromatin fiber results in stronger compartmental segregation [182].

Using (6), χ can be calculated for *any* pair of clutches *e.g*., a HP1α/β/γ-containing clutch (H3K9me3-marked nucleosomes) vs a euchromatic clutch; a Pc-containing clutch (H3K27me3-marked nucleosomes) vs a euchromatic clutch; a HP1 β-containing clutch vs a HP1 γ-containing clutch, and any other pair-wise combinations. This is analogous to the approach for BCPs, where it is the degree of incompatibility between monomers (the “*unit of incompatibility*”) that determines the magnitude and sign of χ (Box 2 and equations (2) to (4)). Here, χ_HC_ specifies the degree of incompatibility between a heterochromatin-*like* clutch and a euchromatic clutch. The magnitude of χ will indicate whether a heterochromatin-*like* domain/complex consisting of repeating units of such clutches has a tendency to micro-phase separate from flanking euchromatin consisting of euchromatic clutches. The degree of phase separation of the domain/complex from the flanking euchromatin is specified by the segregation product χN, where N is the number of repeating units, that is: (i) the number of heterochromatin-*like* clutches in a heterochromatic-*like* domain/complex and, (ii) the number of Pc-containing clutches in Pc-G domain/complex. The magnitude for χN will determine the character of the micro-phase separation actually observed (*i.e*., wavy, “liquid-like” or discrete, sharp interfaces) and this is our subject in the next section.

## 9. χN and the order-disorder transition in relation to heterochromatin-*like* and Pc-G domains /complexes

The sign and magnitude of χ specifies the *degree of incompatibility* between dissimilar monomers that make up the blocks in a symmetrical di-BCP and indicates whether the di-BCP has a tendency to micro-phase separate. The *degree of phase separation* of a symmetric di-BCP is specified by the segregation product χN, where N is the number of repeat units (monomers) that make up the polymer chain [175]. The seminal study that calculated the value of χN at which a bulk di-BCP would phase separate is that of Leibler [211] using self-consistent field theory (SCFT; [212]), where it was predicted that phase separation takes place when χN is around 10.5. This was confirmed experimentally [151]. Given that phase separation of a di-BCP takes place at χN ~10.5 when χN ≈10 there is a delicate balance between energetic and entropic effects on segregation of the blocks (Box 3). When χN is increased there is a first-order phase transition (analogous to the phase transition from water to ice) that is called the order-disorder transition (ODT; χN_ODT_ ~10.5) where a disordered phase that is entropically favoured yet energetically costly is replaced by a periodic lamellae mesophase (χN ≥10; Box 3). The character of the mesophase is dependent upon the magnitude of χN. When χN is close to or slightly greater than χN_ODT_, i.e., χN ≥10, the interfaces between the blocks are weak and “wavy” having the appearance of liquid-liquid phase separation (see ϕ_A_ vs r_⊥_ for χN ≥10 in Box 3). When χN is much greater than the χN_ODT_, *i.e*., χN>>10, the interfaces are discrete and sharp (see ϕ_A_ vs r_⊥_ for χN>>10 in Box 3). The evolution of micro-phase separation as the magnitude of χN is varied (Box 3) highlights how the interface between microphase separated “blocks” can change from wavy, “liquid-like”, to sharp and discrete depending on the magnitude of χN. This is relevant to the character (degree) of the micro-phase separations observed for heterochromatin-*like* compared Pc-G domains/complexes in the interphase nucleus.

### Box 3: The segregation product χN and the order-to-disorder (ODT) transition of BCPs

The degree of micro-phase separation of bulk (undiluted) diblock copolymers (di-BCPs), where the A and B blocks are incompatible, is determined by the segregation product, χN, where N is the number of repeat units (monomers) that make up the polymer chain. Self-consistent field theory [211] predicts that micro-phase separation takes place when χN~10.5, which has been confirmed experimentally [151]. When the di-BCP is symmetrical (volume fraction *f* = 0.5) and χN>10.5 phase separation takes on the character of lamellae. The reason for this is that the covalent linkages stop the A and B blocks from macro-phase separating: the thermodynamic forces driving separation are counterbalanced by entropic forces from the covalent linkage. The resulting chain elasticity keeps the dissimilar A and B portions of the di-BCP apart. As a consequence, symmetric di-BCPs adopt extended configurations seen as lamellae that are in length scales similar to the molecular dimensions of the di-BCPs themselves (0.05-0.1μm).

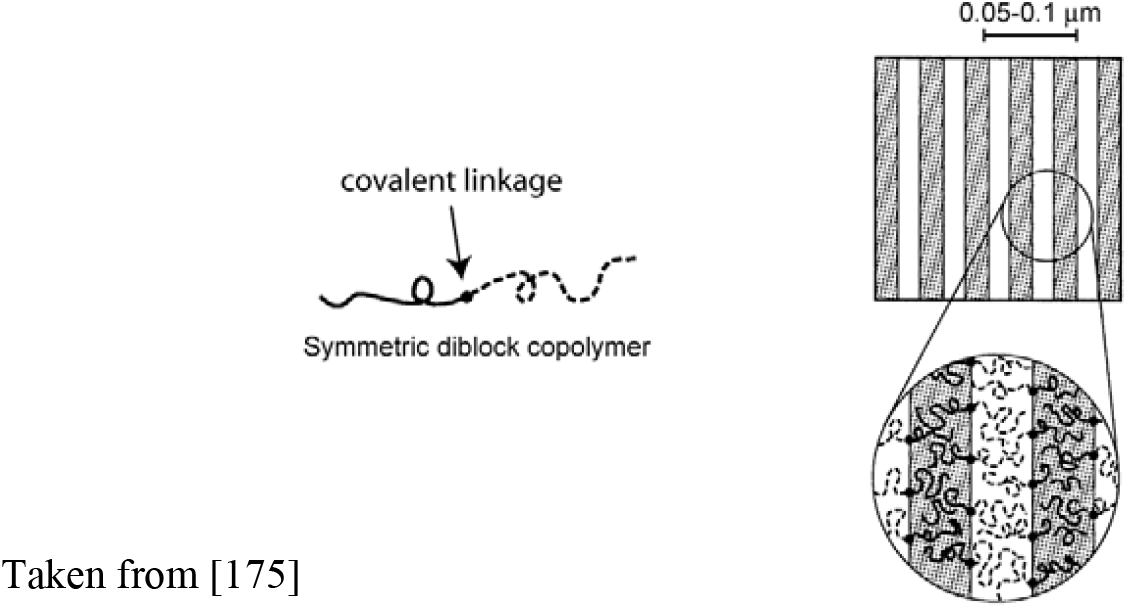

When χN<<10 for symmetric di-BCPs entropic factors dominate and they possess a homogeneous composition profile when plotting composition, ϕ_A_ versus the ensemble average of the bond vectors <r>. When χ or N are increased so that χN ≈10 there are local fluctuations in composition owing to small variations in system entropy (~N^−1^) or energy (~χ) and this results in disordered states (χN ≤10). As χN is increased further there is an orderdisorder transition (ODT; χN_ODT_) where the disordered microstructure is replaced by a periodic lamellae mesophase (χN ≥10) albeit weak A-B interactions still occur and, as consequence, the interfaces are weak and “wavy” (see ϕ_A_ vs r_⊥_ for χN ≥10).

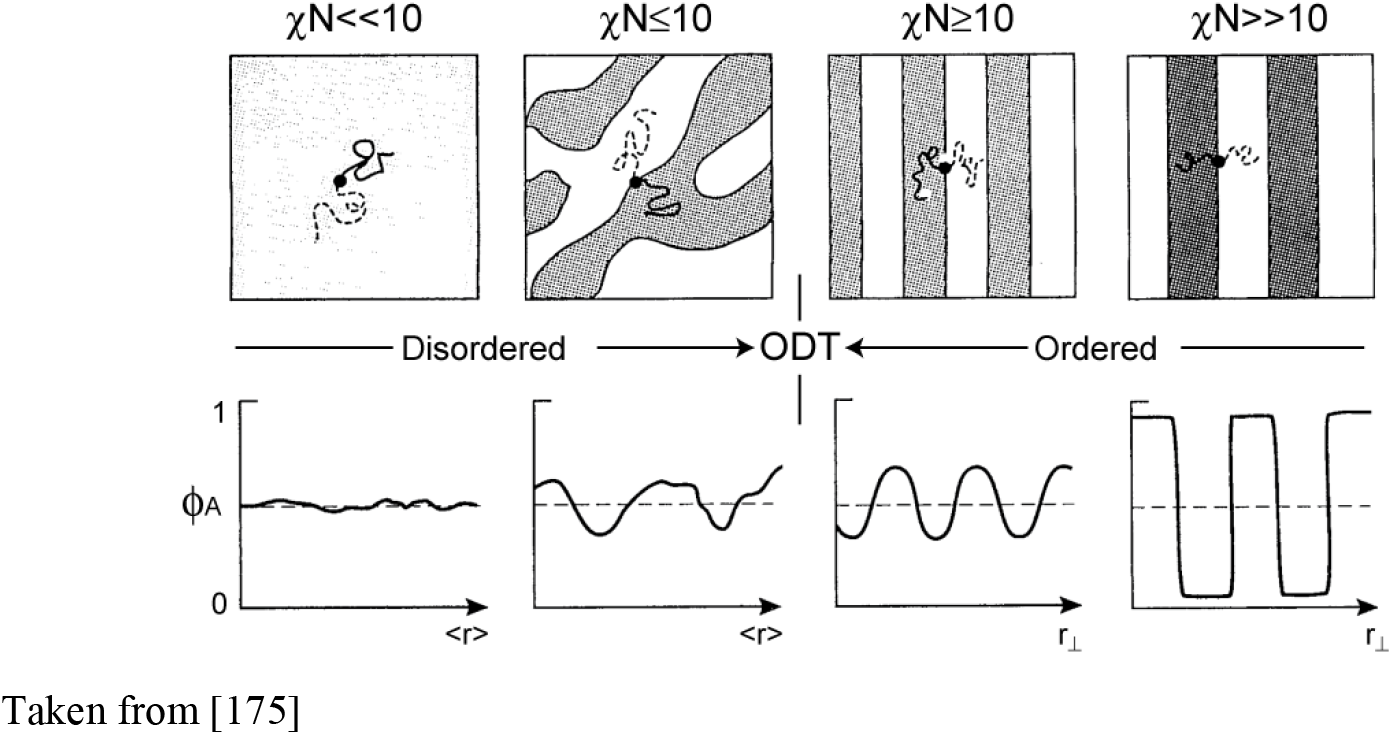

At the limit when χN>>10 energetic factors prevail and a strongly segregated lamellae pattern is observed where interfaces become narrow and micro-domains are composed of pure A or B and the ensemble bond vectors, r, are essentially perpendicular to the periodic mesophase. The evolution of the structures as the product χN increases shows that phase separation close to the ODT (χN≥10) gives rise to interfaces that are weak, wavy, almost liquid-*like*, while much greater than the ODT (χN>>10) interfaces are narrow and sharp. The relationship between χN, χN_ODT_ and degree of phase separation defined analytically with BCPs may provide insight into the character (degree) of the phase separation observed with heterochromatin-*like* domain/complexes compared to that seen with Pc-G domain/complexes. An important caveat is the domains/complexes are unlikely to behave like simple flexible chains governed by Gaussian statistics as assumed for bulk BCPs [151,211]. Because of this, heterochromatin-*like* and Pc-G domains/complexes in the interphase nucleus will have their own values for order-disorder transition, χN_ODT_HC_ and χN_ODT_PC_ respectively.

The original SCFT approach to micro-phase separation of bulk BCPs treats polymers as flexible chains that coarse-grain as Gaussian random walks at sufficiently large (infinite) length scales [151,211]. However, the assumption of random-walk conformations of chains does not accurately describe the behaviour of domains/complexes in the interphase nucleus, where the majority of complexes in mouse and man are small, in the range of 10-30kb (Table S1) albeit domains can be over 1Mb in size (Table 1). A more accurate description is that HP1-containing heterochromatin fibers behave like semi-flexible worm-like chains [185,213]. Semi-flexibility would mean that domains/complexes have their own values for χN_ODT_ distinct from ~10.5 for bulk di-BCPs that behave as flexible chains governed by Gaussian statistics [151,214]. Accordingly, for the discussion below, the values for the order-disorder transition of heterochromatin-*like* and Pc-G domains/complexes flanked by euchromatin are termed χN_ODT_HC_ and χN_ODT_PC_ respectively.

A bulk symmetric di-BCP is a simple one component system where segregation can be described analytically [151,211; Box 3], whereas segregation of heterochromatin-*like* domains/complexes involves a multicomponent system and takes place in the context of a concentrated solution of chromatin fibers where other domains/complexes roam, to a lesser or greater extent, the nuclear space as part of those fibers. Segregation of domains/complexes in interphase nuclei will therefore involve a different physics to that which has been described for bulk symmetrical di-BCPs [151,211; Box 3]. Segregation of domains/complexes takes place in two stages: micro-phase separation then segregation. The magnitude of χ_HC_ – quantified by equation (6) - will indicate the tendency of a heterochromatin-*like* domain/complex to micro-phase separate. The degree of phase separation of the domain/complex is specified by the product χN_HC_ (where N_HC_ is the number of “clutches” in the heterochromatin-*like* domain/complex). When χN_HC_ is greater than χN_ODT_HC_ the domain/complex will be become enriched owing to “bridging” by HP1 of H3K9me3-marked nucleosomes within and between clutches of the domain/complex (Diagram A in Box 1; top row, “HP1 compaction”) resulting in micro-phase separation from the flanking euchromatin (Diagram B in Box 1). Segregation is mediated by a like-with-like attraction (binding potential) between micro-phase separated heterochromatin-*like* domain/complexes that occurs in *cis* or *trans* (Diagrams B and C in Box 1). The binding potential results from an entropic effect. Since the heterochromatin-*like* domains/complexes have already given up entropy as a result of compaction (**S_COMP_** in equation (6)), they have less entropy to lose when they come into contact with each other compared to contact with euchromatic segments of chromatin fibers [213]. Once in contact, HP1 proteins can “bridge” between domains/complexes, forming inter-fiber bridges, which will stabilise far *cis*- and *trans*-contacts that emerge as ECDs is Hi-C experiments (Diagrams B and C in Box 1). Notably, work in *Drosophila* and mouse have shown that HP1 proteins from tightly localised domains in interphase nuclei that cannot be explained in terms of “liquid droplets” that form as a result of liquid-liquid phase separation [41,180,215]. Instead, the character (degree) of this phase separation is consistent with χN_HC_>>χN_ODT_HC_, which would result in sharp interfaces between heterochromatin-*like* domains/complexes and euchromatin, reminiscent of what is observed for di-BCPs when χN is much greater than the χN_ODT_ (χN >> χN_ODT_; see ϕ_A_ vs r_⊥_ for χN>>10 in Box 3). Equation (6) also shows how a value for χ_PC_ for a Pc-containing clutch can be quantified. The magnitude of χ_PC_ will indicate the tendency of a Pc-G domain/complex to micro-phase separate; the degree of micro-phase separation will be given by χN_PC_ (where N_PC_ is the number of “clutches” in the Pc-G domain/complex). Notably, the character of phase separated Pc bodies that contain Pc-G domains/complexes has been reported to be “liquid-like” [216,217]. It is tempting to speculate that this is because χN_PC_ ≥ χN_ODT_PC_ resulting in wavy, liquid-like, interfaces between the Pc-G domains/complexes and euchromatin. This would be similar to what is observed for di-BCPs when χN is close to χN_ODT_ (χN ≥ χN_ODT_; see ϕ_A_ vs r_⊥_ for χN ≥10 in Box 3).

The different degrees of phase separation observed with heterochromatin-*like* domains/complexes compared to Pc-G domains/complexes may reflect the finding that the binding affinity of HP1 for H3K9me3 is much higher than the affinity of Pc CBX homologues for H3K27me3 [218]. Given equation (6) the difference in binding affinity would likely give a value of χ for a heterochromatin-*like* clutch that is greater than that for a Pc-G clutch (χ_HC_ >χ_PC_). This would in turn determine the magnitude of χN_HC_ and χN_PC_ and thereby the character (degree) of phase separation (wavy, “liquid-like” vs sharp interfaces) of the respective domains/complexes.

## 10. Conclusions and perspectives

The hallmarks of constitutive heterochromatin, HP1 and H3K9me3, were present in the common ancestor of fission yeast and man around one billion years ago [16,17]. Both are enriched at canonical sites of constitutive heterochromatin, the peri-centric regions, (sub-) telomeres and (peri-) nucleolar organisers [18–25] and their function at these sites has been the subject of numerous studies in different organisms [204,219]. Less well studied are heterochromatin-*like* domains/complexes that share the hallmarks, HP1 and H3K9me3, with constitutive heterochromatin but lie *outside* the canonical constitutively heterochromatic territories [57]. Our survey shows that domains/complexes are likely to be present in most eukaryotic genomes (Figure 2; Table 1). One of the best characterised heterochromatin-*like* complexes is that which encompasses the 20kb donor mating type region in fission yeast [220], indicating that heterochromatin-*like* domains/complexes represent an ancient mechanism for epigenetically regulating chromatin-templated processes in euchromatic regions of the genome outside canonical constitutive heterochromatin [16].

Heterochromatin-*like* domains/complexes have expanded in mammals where the number in the human genome is, conservatively, around 163-855 heterochromatin-*like* domains (>0.1Mb) and around 18853-32292 complexes (<0.1Mb) (Table 1). How heterochromatin-like domains/complexes domains are nucleated and assembled at specific sites in the genome has been reviewed recently using as an exemplar the B4 sub-compartment that is generated by far *cis*-contacts between the KRAB-ZNF heterochromatin-like domains [41,62]. Briefly, the KRAB-ZNF heterochromatin-like domains (up to 4Mb) are nucleated by smaller complexes (~6kb) assembled at specific sites within the domains by sequence specific KRAB-ZFPs, where the density of nucleation sites is around 30 nucleosomes per 200 nucleosomes [41]. Targeting of nucleation sites by KRAB-ZFPs may be a common mechanism for assembly of larger domains. A recent survey of 222 of the 350 human KRAB-ZFPs showed that the number of genomic sites bound per protein ranged from more than 10,000 to around 15 [221], indicating that KRAB-ZFP-directed heterochromatin-*like* complexes could nucleate larger domains/complexes at many sites within the genome. A majority, 159 of the 222, of KRAB-ZFPs were found to bind at least one type of transposable element (TE) and their binding to TEs is thought assemble local heterochromatin-*like* complexes that repress transposon expression [221,222]. It is common for TE regulatory sequences to undergo exaptation and take on new functions [222,223]. It is tempting to speculate that a number of TE-specific KRAB-ZFP binding sites may have been subject to exaptation and now act as nucleation sites that assemble larger heterochromatin-*like* domains. Other genomic sites include imprinted gDMRs where heterochromatin-*like* complexes are assembled through binding of ZFP57/445 to their methylation-sensitive recognition sequences. Assembly of complexes at imprinted gDMRs preserves parent-of-origin-specific 5mC against the global DNA demethylation that characterises pre-implantation development in mouse and man (Figure 3) [95,116]. In addition to the heterochromatin-like domains/complexes, the related Pc-G domains/complexes number in the region of 2000 in mouse and human genomes [148]. Both heterochromatin-*like* and Pc-G domains are probably involved in regulating the phylotypic progression (Figure 4). As explained (Figure 4), during the phylotypic progression segregation of heterochromatin-*like* domains/complexes likely contribute cell-type-specific contact enrichments to ECDs that safeguard cellular identity, while Pc-containing domains/complexes contribute position-specific contact enrichments. ECDs represent the “epigenetic” component of cellular identity (Diagram C in Box 1; [41]).

Another aspect of cellular identity is age. In this context, it will be of interest to investigate the relationship between ECDs (Box 1) with the “epigenetic clock” developed by [224]. For humans, the “clock” is based on age dependent changes in DNA methylation at 353 ‘clock’ CpGs, where methylation of 193 of the 353 CpGs increase with age while the remaining160 CpGs decrease with age. The epigenetic clock has a high ticking rate until adulthood (~20 years), after which it slows to a constant, steady, ticking rate that can be used to predict the age (or epigenetic age, eAge) of multiple tissues with a median error of 3.6 years [224]). It is known that HP1 proteins interact with DNA methyltransferases [122] and, as explained, HP1proteins are associated with permissive as well as repressive chromosomal regions [64,65] indicating that domains/complexes could regulate both the increasing (193 CpGs) and decreasing (160 CpGs) methylation changes observed with the 353 “clock” CpGs. This remains to be tested.

A key question concerns the contribution of DNA methylation to the generation of ECDs. DNA methylation is an essential epigenetic mechanism that regulates the cell-to-cell inheritance of gene repression patterns [225]. Its role in folding the genome in the interphase nucleus and the emergence ECDs detected in Hi-C maps may be indirect through an effect on the distribution of H3K27me3. DNA hypomethylation can affect the distribution H3K27me3 in the genome [226] most likely owing to targeting of PRC2 and PRC1 to CGIs [227,228] and the generalized affinity of Polycomb repressive complexes for chromatin [229]. Notably, changing the distribution of H3K27me3 by DNA hypo-methylation affects compartmentalisation as detected in Hi-C maps; introduction of DNA methyltransferase activity reconstitutes both H3K27me3 distribution and the Hi-C maps [230].

BCP theory has been used to provide plausible explanations for changes in compartmentalisation seen in Hi-C maps when chromatin-associated cohesin activity is experimentally manipulated [41,156] (Figure 5). Implicit in these explanations is there is a value of χ for heterochromatin-*like* domains/complexes flanked by euchromatin. A recent attempt at determining the degree of incompatibility between A- and B-type homo-polymers (euchromatin vs heterochromatin) estimated χ as 0.03±0.01/nucleosome [182]. This value was thought crude because it was calculated based on phase separation of bulk homo-polymers [178], contrary to the known nuclear environment of a concentrated solution of chromatin “polymer” fibers. We suspect that the problem lies at a more fundamental level. As explained, mean field lattice theory [154,155] used to define χ, assumes the *unit of incompatibility* to be a monomer (Box 2; equations (2) to (4)). Similarly, the seminal exposition on scaling concepts in polymer physics [178] used to estimate χ above [182] subsumes the monomer into a statistical segment (persistence length, *l_p_*); the fundamental unit nevertheless remains the monomer. Instead, we suggest that the *unit of incompatibility* for chromatin is a larger more dynamic structure that probably represents the secondary structure of chromatin, namely the oligo-nucleosomal “clutch”, where a clutch contains from 2 to 10 nucleosomes (see section 8 for details; Figure 6). In our “clutch” model (Section 8) incompatibility is due not to the epigenetic modifications of the nucleosomes *per se*, rather incompatibility is owing to proteins that “bridge” nucleosomes within a clutch, examples of which are HP1 and Polycomb CBX homologues that bridge H3K9me3- and H3K27me3-marked nucleosomes respectively (Diagram A in Box 1). Our approach (Section 8; Figure 6; equation (6)) could be used to quantify χ for both a heterochromatin-*like* “clutch” and a Pc-G “clutch”. For large domains/complexes consisting of N number of clutches the magnitude of χN will determine the character (degree) of phase separation (section 9 and Box 3 for details). Each type of domain/complex has its own value for the order-disorder transition (χN_ODT_) and when χN >> χN_ODT_ the interfaces between the domain/complex and euchromatin will be sharp and well-defined as observed for heterochromatin-*like* complexes [41,180,215]. Where χN ≥ χN_ODT_ the interfaces between the domain/complex and euchromatin will be wavy, “liquid-like”, as observed for Pc bodies that contain Pc-G domains/complexes [216,217].

The mesoscale organisation of chromatin as oligo-nucleosomal “clutches” containing 2 to 10 nucleosomes with a variable zig-zag organisation of nucleosomes appears to be a ubiquitous motif of interphase chromatin [188]. What causes this organisation to emerge from the “sea of nucleosomes” [188,231,232] that characterises interphase chromatin is not known. Given our model (Section 8) and equation (6) the mesoscale organisation could arise from (transient) “bridging” of nucleosomes by proteins that possess two (or more) chromatin-binding motifs [233,234]. The sum of competing free energy contributions (for a heterochromatin-*like* clutch this would be (**H_CD-H3K9me_**)_HC_ and (**H_CSD-CSD_**)_HC_ in equation (6)) and potential energies (**H_COMP_** and **H_TL_** in equation (6)) could generate, in the dynamic environment of the interphase nucleus, oligo-nucleosomal “clutches” that vary from 2 to 10 nucleosomes. Of note is that a simple application of equation (6) to constitutive heterochromatin, where both HP1 and H3K9me3-marked chromatin fibers are highly-enriched [29], predicts the formation of larger clutches with more pronounced zigzag geometry. This is, in fact, what is observed [189,194].

## 11. Coda

- The oligo-nucleosomal “clutch” is the “*unit of incompatibility*” of chromatin.
- When a domain/complex assembles from “clutches” consisting of “bridged” nucleosomes with zig-zag geometry it will have a tendency, specified by χ, for micro-phase separation from flanking euchromatin.
- A *qualitative* prediction of the degree of micro-phase separation (wavy vs sharp interfaces) is specified by the magnitude of χN compared to χN_ODT_ for a given domain/complex.
- Segregation of micro-phase separated domains/complexes facilitates folding of the genome into the confines of the nucleus while retaining an environment that enables (regulation of) chromatin-templated processes.

## Abbreviations

5mC: 5-methylcytosine
Å: Angstrom
*α*: segment polarizability
α: coefficient for enthalpy
ADD: ATRX-DNMT3-DNMT3L domain
A-P: anterior-posterior
ATRX: Alpha Thalassemia/Mental Retardation Syndrome X-Linked
β: coefficient for excess entropy
BCP: block copolymer
bp: base pair
CBX: Chromobox homologue
CD: chromodomain
cen3: fission yeast centromere 3
CGIs: CpG islands
ChAHP: CHD4-ADNP-HP1 complex
CpG: CpG di-nucleotide
CSD: chromo shadow domain
CTCF: CCCTC-binding factor
DAXX: Death Domain associated protein
di-BCP: di-block copolymer
DNMT1: maintenance DNA methyltransferase 1
ε_AB_, ε_AA_, and ε_BB_: contact energies per repeat unit of A–B, A–A, and B–B, respectively
ECD: epigenetic compartmental domain
*ε_ij_*: contact energy
ε_ps_, ε_pp_, and ε_ss_: contact energies of polymer–solvent, polymer–polymer, and solvent–solvent, respectively
Eso1: SMC3 acetyltransferase
ES: embryonic stem
FRAP: fluorescence recovery after photo-bleaching
gDMRs: germline differentially methylated regions
*f_A_*: volume fraction of A
*f_B_*: volume fraction of B
H3K4me3: *tri*-methylated lysine 4 on histone H3
H3K9me2: *di*-methylated lysine 9 on histone H3
H3K9me3: *tri*-methylated lysine 9 on histone H3
H3K27me3: *tri*-methylated lysine 27 on histone H3
H4K16: lysine 16 of histone H4
H4K20me3: *tri*-methylated lysine 20 on histone H4
HMTases: histone methyltransferases
HP1: Heterochromatin Protein 1
*Hox*: homeobox
HR: hinge region
ICM: inner cellular mass
KAP1: KRAB-associated protein 1
kb: kilobases
*k_B_*: Boltzmann’s constant
*k_B_T*: thermal energy
kDa: kiloDalton
*I*: first ionization potential
*i* and *j* segments: “blocks” of a di-BCP each consisting of (dissimilar) monomer repeating units
KRAB-ZNPs: Krüppel-associated box (KRAB) domain zinc-finger proteins
KRAB-ZNF: KRAB domain-zinc finger
LEF: loop-extruding factor
Mb: Mega-bases
ncRNA: non-coding RNA
*N* = *N_A_* + *N_B_*: total degree of polymerization of AB di-BCP
NORs: nucleolar organiser regions
ODT: order-disorder transition
Pc-G: Polycomb-Group consisting of H3K27me3, PRC1 and PRC2 complexes
Pds5A/B: sister chromatin cohesin protein 5A/B
*l_p_*: persistence length
PEV: position-effect variegation
PI-b-PS: polyisoprene-block-polystyrene
PRC1: Polycomb-repressive complex
PRC2: Polycomb repressive complex 2
RICC-Seq: Radiation-induced spatially correlated cleavage of DNA with sequencing
PxVxL: Proline/Any/Valine/Any/Leucine penta-peptide motif
*r_ij_*: segment-segment separation
ϕ_A_: composition of A
<r>: ensemble average of bond vectors
r_⊥_: ensemble bond vectors perpendicular to periodic mesophase
SETDB1: SET Domain Bifurcated 1 K9H3 HMTase
Sir: silent information regulator
SCC1: cohesin subunit Scc1
SCM3: cohesin subunit SMC3
SUMO2: Small ubiquitin-related modifier 2
Suv39h1/2: mammalian suvar K9H3 HMTase 1 and 2
Swi6^HP1^: HP1 homologue swi6p from fission yeast
T: temperature
TAD: topologically associated domain
TEs: transposable elements
Tet: Ten-eleven translocation dioxygenase
UBE2i: Ubiquitin conjugating enzyme 2i
WAPL: WAPL cohesin release factor
χ: The Flory-Huggins parameter
χ_HC_: specifies the degree of incompatibility between a heterochromatin-like “clutch” and a euchromatic “clutch”
χN: segregation product that specifies the degree of phase separation
χN_ODT_HC_: magnitude of segregation product at the order-disorder transition for a heterochromatin-like domain/complex
χN_ODT_PC_: magnitude of segregation product at the order-disorder transition for a Pc-G domain/complex.

## Availability of data and materials

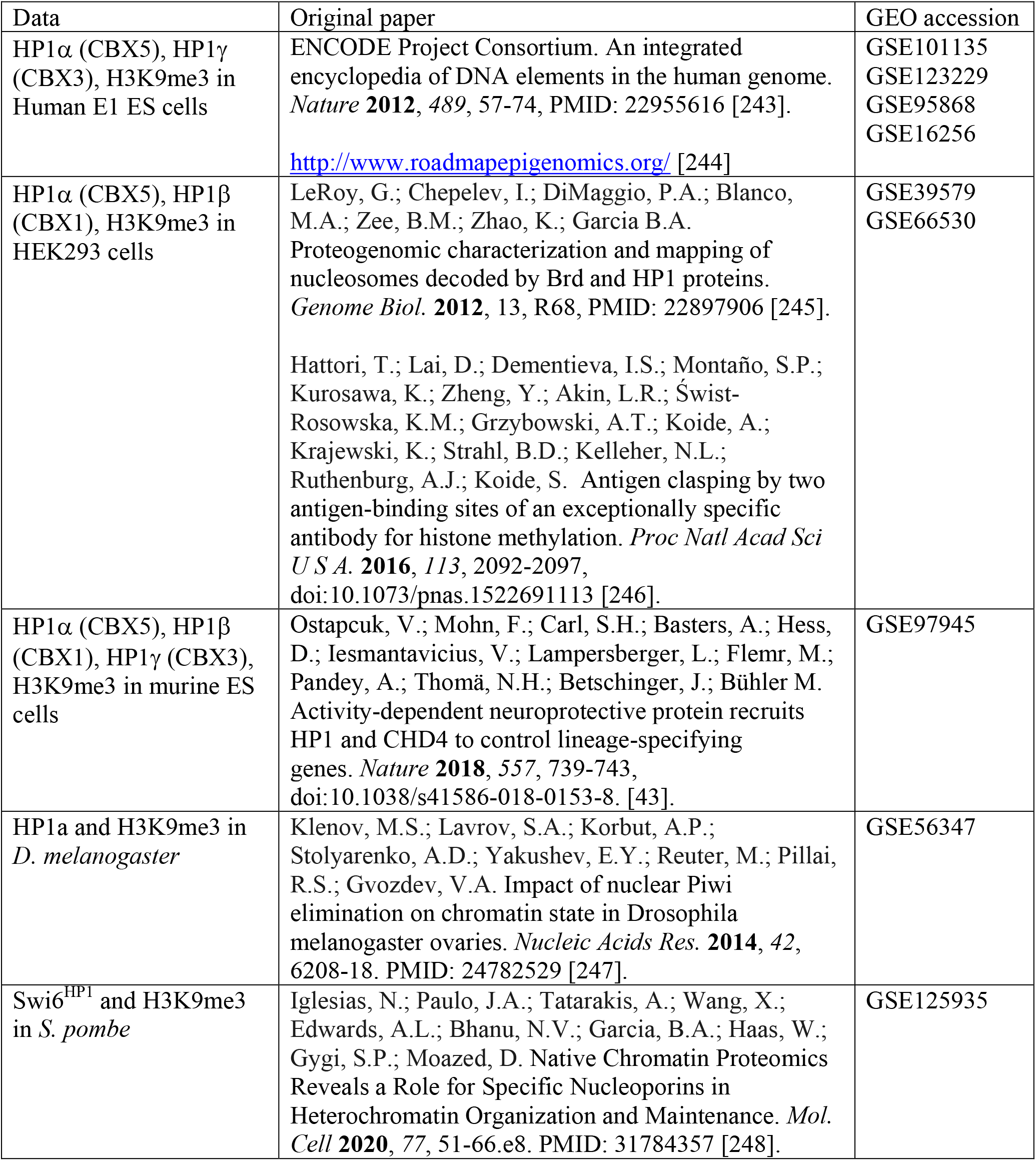

### Data analysis

ChIP-seq raw data were trimmed for quality using Trimmomatic SE and aligned to GRCh38 (Ensembl91, Human), GRCm38 (Ensembl95, Mouse), BDGP6 (Ensembl95, *D. melanogaster*) and ASM294 (*S. pombe*) using Bowtie2 [249]. Assemblies in which constitutive heterochromatin regions were absent or unmapped were used; we also excluded constitutively heterochromatic regions based on the ENCODE blacklist [250]. ChIP peak calling and fold enrichment analysis was performed in MACS2 tool using–*broad* parameter [251]. Correlation analysis of fold enrichment of ChIP-seq profiles was performed using multiBigwigSummary from deepTools and R package *ggpubr* [252]. Genome-wide annotation of heterochromatin marks enriched regions was performed with ChromHMM tool [253].

## Data accessibility

See the Availability of data and materials section above.

## Author contributions

PBS conceived of the synthesis presented, wrote the paper and drafted the figures. SNB drew all figures except for figure 2. PPL undertook the bioinformatic analyses and assembled figure 2. All authors made comments on the manuscript and approved the final version.

## Acknowledgements

We are grateful to Andrew Newman for advice on bioinformatics and many helpful discussions. We thank Andrew Newman, Jafar Sharif and Professor FS Bates for comments on the manuscript.

## Footnotes

1 H3K9me2/3 will be termed H3K9me3 [14].

2 This equation, as given in [175], contained a numerical error, where the coefficient should be 3/8 (as it is here). The corrected equation (4) above was graciously provided by Professor FS Bates.

