## Supplemental Table 1 for "Biology and physics of heterochromatin-*like* domains/complexes"

|  | *H. sapiens* \| HP1α,HP1γ,H3K9me3\| H1 ES Cells | | *H. sapiens* \| HP1α,HP1β,H3K9me3\| 293T HEK Cells | | *M. musculus* \|  HP1α,HP1β,HP1γ,H3K9me3 \|  ES cells | |
| --- | --- | --- | --- | --- | --- | --- |
| Size, kb | N | % | N | % | N | % |
| 100-90 | 30 | 0.16 | 229 | 0.71 | 223 | 2.18 |
| 90-80 | 24 | 0.13 | 328 | 1.02 | 271 | 2.65 |
| 80-70 | 35 | 0.19 | 477 | 1.48 | 364 | 3.56 |
| 70-60 | 85 | 0.45 | 667 | 2.07 | 566 | 5.53 |
| 60-50 | 154 | 0.82 | 1043 | 3.23 | 725 | 7.09 |
| 50-40 | 403 | 2.14 | 1845 | 5.71 | 1119 | 10.94 |
| 40-30 | 1118 | 5.93 | 3455 | 10.70 | 1583 | 15.48 |
| 30-20 | 3356 | 17.80 | 7014 | 21.72 | 2427 | 23.73 |
| 20-10 | 13648 | 72.39 | 17234 | 53.37 | 2949 | 28.84 |
| ***Total*** | ***18853*** |  | ***32292*** |  | ***10227*** |  |

**Table S1: Size range of heterochromatin*-like* complexes in man (H1 ES cells and 293T cells) and mouse (ES cells)**
